# The direction and timing of theta and alpha traveling waves modulate human memory processing

**DOI:** 10.1101/2022.02.07.479466

**Authors:** Uma R. Mohan, Honghui Zhang, Joshua Jacobs

**Affiliations:** Department of Biomedical Engineering, Columbia University; Amazon Corporation, Seattle, Washington; Department of Neurological Surgery, Columbia University

## Abstract

To support a range of behaviors, the brain must flexibly coordinate neural activity across widespread brain regions. One potential mechanism for this coordination is a traveling wave, in which a neural oscillation propagates across the brain while organizing the order and timing of activity across regions^1,2^. Although traveling waves are present across the brain in various species^3–5^, their potential functional relevance remained unknown. Here, using rare direct human brain recordings, we demonstrate two novel functional roles for traveling waves of theta- and alpha-band (2–13 Hz) oscillations in the cortex. First, traveling waves propagate in different directions during separate cognitive processes. In episodic memory, traveling waves tended to propagate in posterior-to-anterior and anterior-to-posterior directions, respectively, during encoding and retrieval. Second, traveling waves are informative about the timing of behavior, with the phase of ongoing traveling waves indicating when subjects would retrieve memories. Because traveling waves of oscillations correspond to local neuronal spiking, these patterns indicate that rhythmic pulses of activity move across the brain with different directions and timing for separate behaviors. More broadly, our results suggest a fundamental role for traveling waves and oscillations in dynamically coordinating neural connectivity, by flexibly organizing the timing and directionality of network interactions across the cortex to support cognition and behavior.

## Introduction

The brain supports a diverse range of behaviors, which requires the coordination of neural activity between different sets of regions. How does the brain support this flexibility? One potential mechanism for flexibly organizing large-scale neuronal activity is a traveling wave (TW), which is a neuronal oscillation that propagates across the cortex^1,2^. TWs are widespread in the brain, appearing across multiple regions in animals^6–8^ and humans^4,9,10^, at both small^11–14^ and large^15–19^ scales. Because TWs correlate with local neuronal activity, their spatiotemporal organization indicates which cortical regions are active and where activity is propagating at each moment^5,15^. Further, due to TWs’ ability to rapidly reorganize^20^, they may support the brain’s ability to dynamically adapt its processes to meet changing environmental demands^21,22^. However, despite these theoretical features and TWs’ widespread prevalence^2,15^, their behavioral importance is unknown. Thus, our goal here was to identify potential functional roles of TWs in human cognition.

Two key properties of TWs are their propagation direction and timing. As a TW propagates, it reflects a moving wave of rhythmic neuronal activity, which causes neurons across neighboring cortical regions to activate in different orders according to the direction of wave propagation^9,23^. Thus, a TW’s direction of propagation may indicate the sequence of activity across neighboring cortical regions, with direction changes signaling a reorganization of the underlying neural connectivity and computation. In this way, separate neural processes and their associated behaviors might be reflected by TWs propagating in different directions^24,25^.

A complementary feature of a TW is its timing, measured via its phase. The instantaneous phase of a TW indicates the positions of the wave’s peaks and troughs along the cortex. In earlier studies, the phase of neural oscillations in many regions correlated with the functional state of the local neuronal network^26–30^, with specific phases indicating different processing states, such as the level of sensitivity to new inputs^5^. Therefore, more generally, we hypothesized that as a TW propagates through a region of the human cortex, its instantaneous phase at a particular region would be informative about the state of local neuronal processing.

Together, via changes in direction and phase, TWs may provide a mechanism to flexibly organize large-scale brain activity to support different behavioral processes. We examined this hypothesis in the domain of human memory, by measuring human TWs directly from neurosurgical patients performing memory tasks. We found that the propagation direction and timing of the brain’s ongoing TWs changed in relation to memory encoding and recall processes. These results demonstrate that different human cognitive processes are supported by large-scale patterns of oscillations that are TWs, with their propagation direction and timing indicating the reorganization of cortical interactions to support behavior.

## Results

### Measuring traveling waves in the human cortex

To examine how the direction and timing of traveling waves (TWs) in the human brain related to cognition, we examined electrocorticographic (ECoG) brain recordings from neurosurgical patients performing memory tasks. The dataset consisted of recordings from 68 subjects performing an episodic-memory task^31^ and 77 patients performing a working-memory task^32^. During these tasks, subjects showed brain oscillations at various frequencies across widespread brain regions, consistent with earlier work^31,33^.

We analyzed these multichannel recordings using spectral analysis and circular statistics to identify the neural oscillations that behaved as TWs and assess their functional role^4,34^. A prerequisite for a brain region to show a TW is that there must be an oscillation at the same frequency across a contiguous region of cortex. Thus, to identify TWs, in each patient we first identified the spatially contiguous clusters of five or more electrodes that simultaneously showed oscillations at similar frequencies, which we refer as “oscillation clusters.” Next, we tested whether each oscillation cluster showed a TW by measuring whether the timing of these oscillations shifted progressively with the position of the electrode within the cluster. To statistically test each cluster for a TW, we measured the instantaneous phase of the oscillation at each electrode and identified consistent phase gradients across neighboring electrodes (see *Methods*). A phase gradient across an oscillation cluster indicates that a TW is present because it means that the cycles of one oscillation are appearing with a progressive delay across neighboring regions of cortex (Fig. 1).

**Figure 1:**
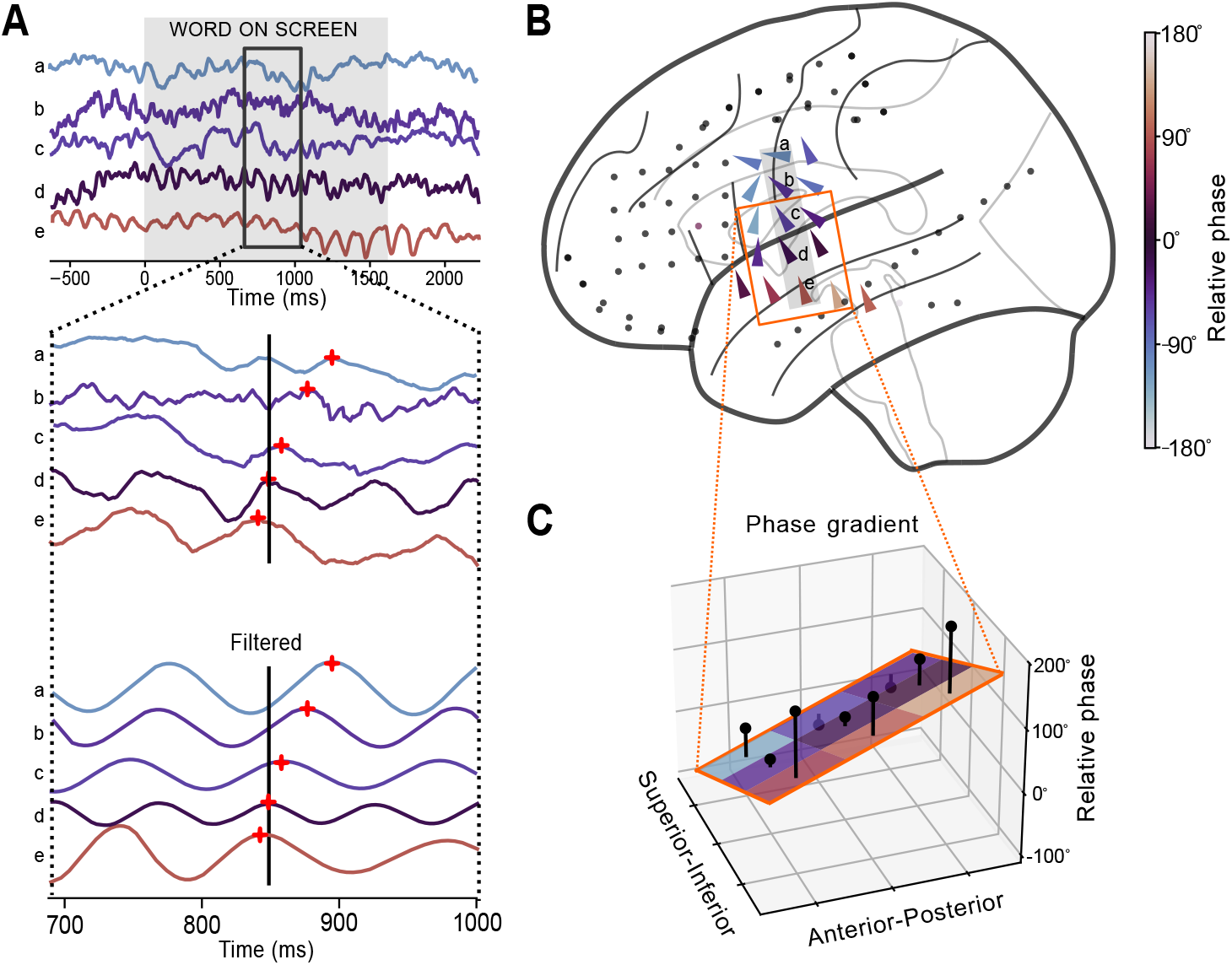
Example traveling wave (TW) at 8.9 Hz in Patient 34’s left hemisphere. **(A)** Recording from one trial of the memory task. Top: raw signal from five selected electrodes. Middle: Expanded view of the signals from the top panel. Bottom: The signals from the middle panel after filtering at 8.9 ± 1.3 Hz. Color indicates relative phase, measured at the time of the black line. **(B)** Brain map indicating the TW in this trial. Arrows indicate, for each electrode, the local propagation direction. Arrow color indicates relative phase at time indicated by black line in A. **(C)** Illustration of the circular–linear regression model for measuring the properties of each TW. This model estimates the spatial phase gradient at each electrode based on the phases from the nearby electrodes’ filtered signals. Black dots indicate the measured phase on each electrode, the plane indicates the model fit, and black lines indicate residuals. The slope of the fitted surface provides an estimate of the TW’s propagation direction and speed.

As in earlier work^4,35^, TWs were widespread in these datasets. We observed significant oscillations and TWs across all brain lobes, in both hemispheres, at frequencies from 2 to 30 Hz. Overall, 72% of electrodes were part of at least one oscillation cluster (Tab. S1). 84% of oscillation clusters exhibited significant TWs (Figs. S1, S2). TWs were prominent during both the episodic and working memory tasks (in 59 of 68 subjects and 64 of 77 subjects, respectively; Tables S2, S3).

Figure 1A illustrates a TW at ∼8.9 Hz that appeared in one trial of the episodic memory task in a patient’s left temporal and frontal cortices. This oscillation was a TW because its individual cycles appeared with a progressive delay across neighboring electrodes. Each cycle of this TW appeared first on ventral electrodes, and then later on anterior–superior electrodes, propagating with a speed of ∼1 m/s. Although the TWs on this cluster varied over time, across trials the propagation of TWs here most often in an anterior–superior direction (Fig. 1B). We measured the propagation of TWs throughout the task using circular statistics (Fig. 1C), which revealed the instantaneous direction, phase, speed, and strength (spatial consistency of the phase gradient). We then tested these features for links to behavior (see *Methods*).

To identify features of TWs that correlated with separate cognitive processes, we examined recordings from an episodic memory task, which previously had revealed brain signals distinguishing distinct stages of memory^33,36^. In each list in this task, subjects viewed a sequence of words and, after a delay, tried to freely recall as many of them as possible (Fig. 2A). On average, subjects successfully recalled 27% of viewed words. Because subjects only remembered a subset of the words in the task, it provided data for us to test whether features of TWs differed according to whether memory encoding was successful.

**Figure 2:**
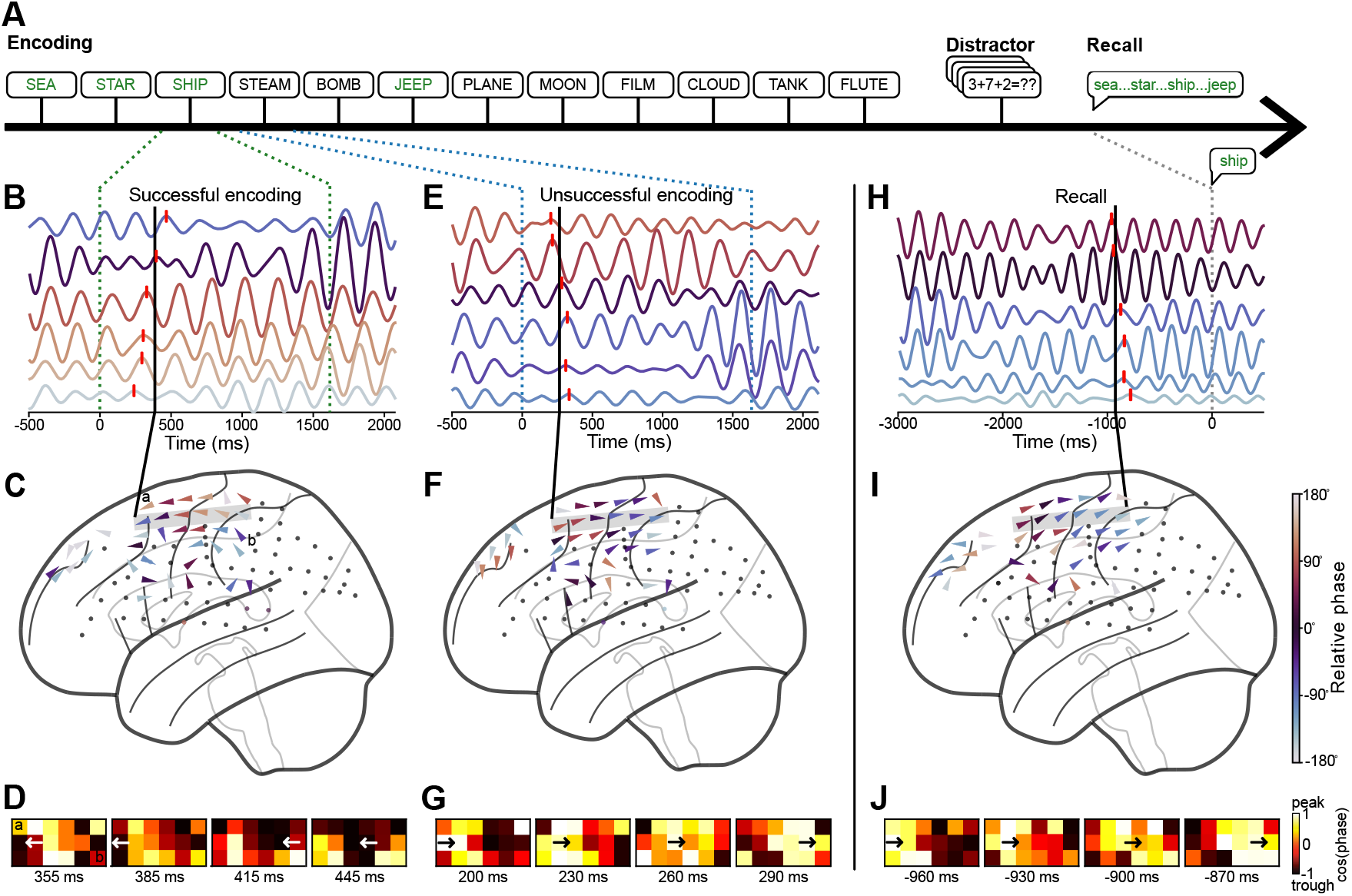
Changes in traveling wave direction across memory encoding and recall. **(A)** Timeline of one trial of the verbal memory task for patient 69. Words colored green were successfully encoded, black words were forgotten. **(B)** Recordings on six electrodes in one trial while the subject successfully encoded the word “SHIP”. Signals were filtered at 4.5 Hz and electrodes were ordered from anterior (top) to posterior (bottom). Red ticks indicate peaks of one oscillation cycle, which illustrates an example TW because there was a progressive shift in the timing of these peaks across electrodes. **(C)** Brain map with arrows showing the direction of TW propagation for the timepoint labeled with the black line in B. **(D)** Topography of this TW’s propagation across a 3 × 6 array of electrodes within the oscillation cluster from panel C (Labels (a) and (b) indicate corresponding electrodes). Each panel indicates the topography of instantaneous phase at one of four sequential time points. **(E–G)** A representative TW measured during unsuccessful memory encoding where the subject viewed the word “STEAM”, with plots analogous to panels B–D. **(H–J)** A representative TW measured prior to the recall of the word “SHIP.”

Figure 2B–J shows data from an oscillation cluster in the frontal lobe of patient 69 with TWs at ∼4.5 Hz during memory encoding and recall. In one trial when the subject viewed a word that they successfully encoded into memory, the electrodes in this cluster showed a TW that propagated in a posterior-to-anterior direction (Fig. 2B–D). Inversely, later in that same list when the subject viewed a different word that they did not successfully encode, there was instead a TW propagating in the opposite, anterior-to-posterior direction (Fig. 2E–G, see also Videos S1, S2). Finally, during recall, before the subject said the name of the remembered word, this oscillation cluster showed a TW propagating in an anterior-to-posterior direction (Fig. 2H–J). This pattern of results—in which the direction of TW propagation shifted according to the current memory process and performance—led us to systematically test the link between different memory processes and TW propagation direction.

### Traveling waves propagate anteriorly during successful memory encoding

We examined the link between TW propagation direction and memory encoding by comparing the properties of the TWs that appeared during the presentations of words that were remembered versus those that were forgotten. A representative example of our results is shown in Figure 3A,B for the same oscillation cluster shown above. Here, when the subject viewed words that they successfully encoded into memory, theta TWs propagated in a posterior-to-anterior direction (*p <* 10^*−*4^, Rayleigh test). When the subject viewed words they did not successfully remember, the TWs here propagated bidirectionally, in a posterior-to-anterior direction on some trials and in an anterior-to-posterior direction on other trials (Fig. 3B, middle). Thus, there was a significant difference in the directions of TW propagation between successful and unsuccessful encoding, with unidirectional posterior-to-anterior propagation for successful memory encoding and bidirectional propagation for unsuccessful encoding (Fig. 3B, *p <* 10^*−*3^, Watson–Williams test). Overall, it was common for clusters to exhibit TWs that propagated bidirectionally (60% of all clusters), by switching over time between propagation in two distinct directions (Figs. S3, S4A, S5; see *Methods*). Overall, subjects who showed bidirectional TW propagation showed a 23% higher rate of successful memory encoding compared to subjects without this pattern (Fig. S4B), indicating that bidirectional TW propagation is a feature of normal cognition.

**Figure 3:**
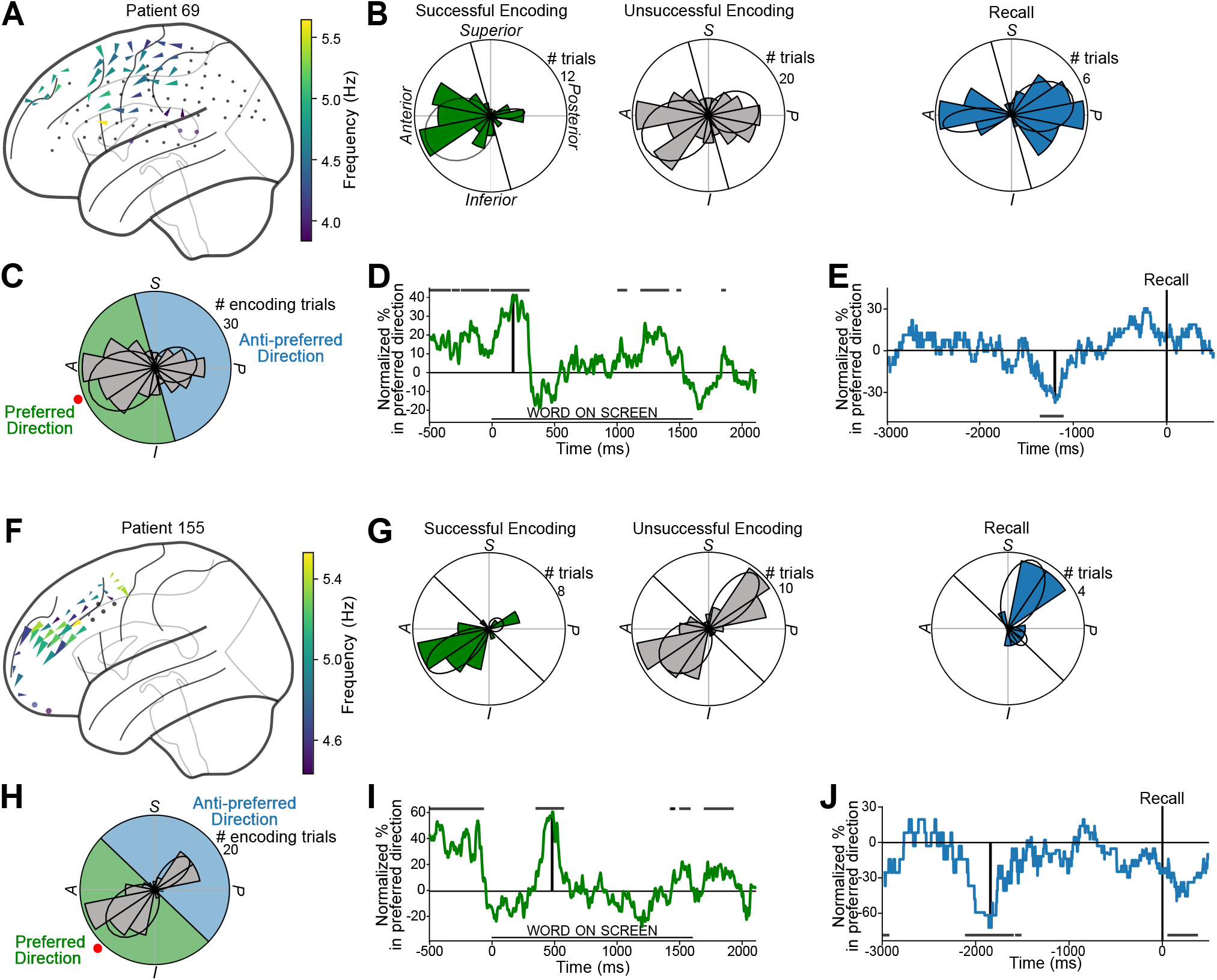
Traveling waves (TWs) vary propagation direction with memory processing. **(A)** Brain map showing the mean direction and frequencies of TWs measured in the left hemisphere of patient 69 during successful memory encoding. Arrows indicate the mean propagation direction for each electrode averaged across trials. Arrow size indicates directional consistency. **(B)** Distribution of TW propagation directions across trials, averaged across the electrodes from panel A, during successful memory encoding (left), unsuccessful encoding (middle), and recall (right). Predominant directional clusters indicated by black ellipses (see Methods). **(C)** Propagation directions of TWs across all encoding trials. The preferred direction is marked with a red dot; green and blue shading indicate the range of angles labeled as preferred and anti-preferred directions, respectively. **(D)** Timecourse of the link between TW propagation direction and memory encoding. Line indicates the difference in the percentages of TWs moving in the preferred direction between trials with successful memory encoding compared to unsuccessful encoding. Vertical black line indicates the time of the maximal difference, which corresponds panels A–C. Horizontal black lines indicates timepoints when directional shifts are statistically significant (permutation test, p < 0.05). **(E)** The link between TW propagation direction and memory recall. The line indicates the normalized percentage of trials propagating in the preferred direction prior to memory recall at time 0. Values are normalized relative to the cluster’s baseline period. Vertical black line indicate the timepoint of the peak anti-preferred propagation (which matches the right panel of B). Horizontal black lines indicate significant timepoints measured by binomal tests. **(F-J)** Same as A–E for patient 155.

We next examined across the entire dataset whether TW propagation direction correlated with memory encoding. We identified the oscillation clusters with bidirectional TW propagation and measured each cluster’s “preferred direction,” which is the propagation direction that was most closely associated with successful memory encoding (Fig. 3C, see *Methods*). We then labeled each timepoint of each trial according to whether the TWs were propagating in the cluster’s preferred or anti-preferred direction. Then, using permutation statistics, we tested the link between propagation direction and whether the subject successfully encoded the viewed word at each timeopint. Using this procedure, in Patient 69 we found a reliable link between memory encoding and TW direction that was strongest 160 ms after word presentation (Fig. 3D; *p <* 0.05, cluster-based correction for multiple comparisions). At this timepoint, when this cluster showed a TW propagating in the preferred direction (i.e., posterior-to-anterior), the subject was 2.2× more likely to remember the word successfully than when the TW was propagating in the anti-preferred direction (35% versus 17% respectively; *p <* 0.01, binomial test, Fig. 3D). Other subjects also showed similar patterns, with significantly better memory encoding when TWs propagated in the preferred direction (Figs. 3F–J, S6).

Consistent with these examples, overall the preferred directions for TWs on individual oscillation clusters were most often posterior-to-anterior (Fig. 4A, top left, *p <* 0.05, Rayleigh test). In contrast, propagation directions for unsuccessful memory encoding shifted and showed increases in anti-preferred and bidirectional propagation (Fig 4A, top right), which was significantly different compared to the distribution of directions during successful encoding (*p <* 0.05, Watson–Williams test).

**Figure 4:**
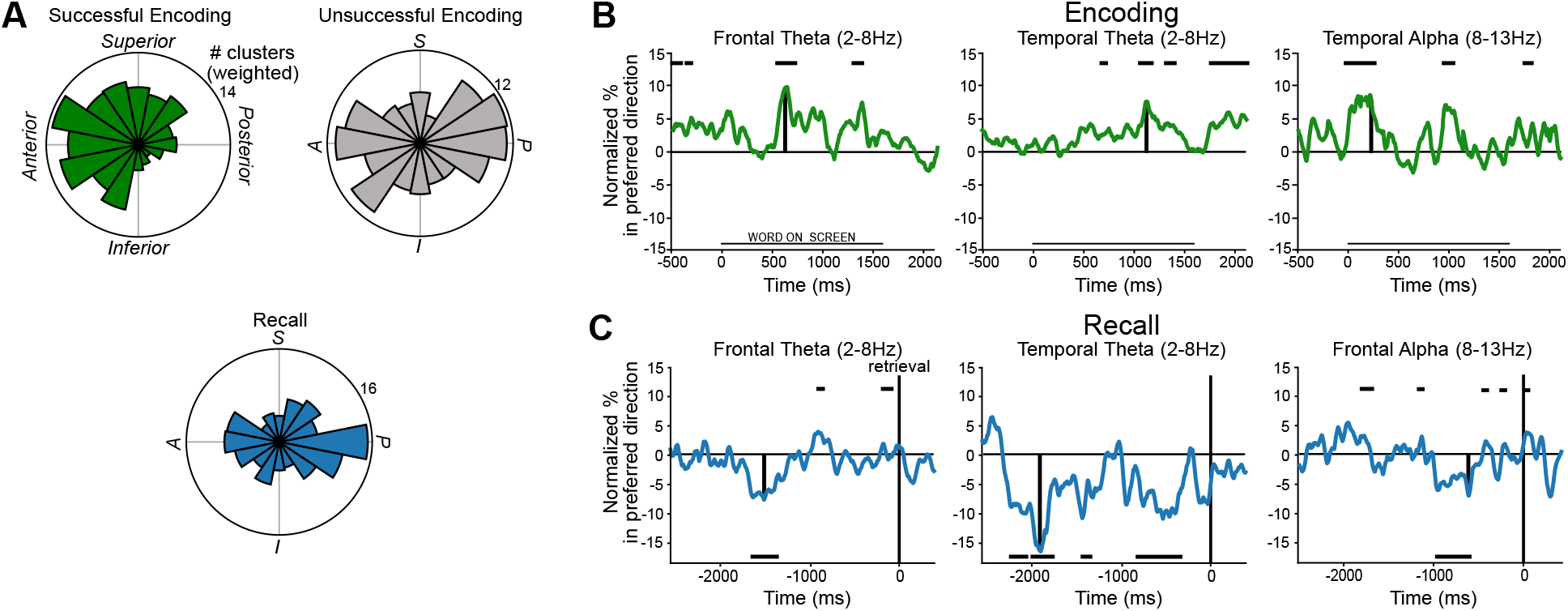
Population analysis of traveling-wave (TW) direction shifts during memory encoding and recall. **(A)**Distribution of clusters’ predominant propagation directions for TWs across all regions and frequencies during memory encoding and recall. **(B)** Timecourses of TW directional shifts during successful and unsuccessful memory encoding. Black vertical lines indicate timepoint of peak propagation in the preferred direction. Horizontal black line indicates statistical significance at p < 0.05 based on permutation testing, with the position at top or bottom of the plot indicating the direction of the effect. **(C)** Timecourses of TW directional shifts prior to memory recall. Black vertical lines indicate times of peak anti-preferred propagation.

Overall, memory-related TWs were widespread. Of the oscillation clusters that showed bidirectional propagation, 69% exhibited a preferred direction that was significantly associated with successful memory encoding (*p*′*s <* 0.05, FDR-corrected binomial tests: see Methods; Fig. S7, Table S1). This link between memory performance and the direction of TW propagation was present at significant levels in the theta band (2–8 Hz) in the frontal and temporal lobes and in the alpha band (8–13 Hz) in the temporal lobe (*p*′*s <* 0.05, binomial tests).

For alpha-band TWs in the temporal lobe, propagation direction significantly correlated with memory encoding even before the word was presented, beginning 24 ms before word presentation and peaking 229 ms after word presentation (p *<* 0.05, cluster-based permutation test, see *Methods*, Fig. 4B, right). At this peak timepoint, subjects showed 1.4× greater memory encoding performance when TWs propagated in the cluster’s preferred direction. Similarly, frontal and temporal-lobe theta-band TWs showed increased propagation in the preferred direction ∼600–1200 ms after word presentation (Fig. 4B, left and middle), and this effect predicted ∼1.8× and ∼1.4× increases in the rate of memory encoding success, respectively (*p <* 10^*−*3^ and *p <* 0.05, permutation tests).

We considered the possibility that this correlation with memory could be more strongly driven by other features of TWs, such as the power of ongoing oscillations, rather than propagation direction specifically. However, we neither found a significant relation between memory encoding and the power of ongoing oscillations nor with the speed or strength of TWs (all *p*′*s >* 0.05; Tab. S4, Fig. S8). Thus, our results indicate that the link between TWs and memory encoding was specific to the direction of propagation.

### Traveling waves propagate posteriorly during memory recall

Immediately before the subject verbally recalls each word, they are actively searching their memory^36,37^. We hypothesized that a different pattern of TWs would be present during this period. To examine the propagation of TWs during memory recall, we examined the same cluster of electrodes (patient 69) during the period prior to the patient speaking aloud the remembered item (Fig. 3E). Here, rather than the posterior-to-anterior propagation that appeared during encoding, instead TWs tended to propagate in the cluster’s anti-preferred, or anterior-to-posterior, direction (Fig. 3B, right). This cluster’s propagation direction during recall was reliably different compared to successful encoding (*p <* 0.05, Kuiper test) and was strongest −1195 ms prior to word recall (Figs. 3E). Thus, the direction of TW propagation on this cluster correlated with the current memory process, switching directions between successful encoding and recall. TWs in other subjects also showed similar patterns (Figs. 3F–J, S6).

Across the dataset, TWs on 40% of the oscillation clusters with bidirectional propagation exhibited a significant pre-recall direction shift. This usually involved increased anterior-to-posterior propagation prior to recall (Fig 4A, bottom, *p <* 0.01, Rayleigh test). Pre-recall direction shifts occurred at significant levels for theta-band TWs in the frontal (7%) and temporal lobes (15%) and for alpha-band TWs in the frontal lobe (7%) (Fig. 4C; all *p*′*s <* 0.05 FDR corrected, binomial tests; Fig. S7, Table S1). Thus, memory recall is associated with theta- and alpha-band TWs in the temporal and frontal lobes that propagate in an anterior-to-posterior direction, which is the opposite direction from memory encoding.

### Traveling wave phase predicts the timing of memory retrieval

In addition to propagation direction, another key characteristic of a TW is phase, which indicates the current location on the cortex of the TW’s peaks and troughs. We thought that the phase of TWs might be relevant for memory because empirical studies showed that the phase of neuronal oscillations predicted perception and attention^5,26–30^ and models described how certain items were represented in memory at specific oscillation phases^38,39^. Building upon this, we hypothesized that the phase of TWs would be informative about the brain’s current state by indicating the specific areas of cortex that most actively support memory retrieval at each moment. The phase of the TW at each location cycles rapidly as the oscillation propagates across the brain. Therefore, to test if TW phase was relevant functionally, we needed a behavioral measure that would be sensitive to the timing of neural processes.

To examine this issue, we examined subjects’ reaction times as they performed the Sternberg working-memory task, which is a paradigm that elicits reliable response timing patterns^32^. We then tested for a relation between TW phase and memory retrieval by comparing each subject’s reaction times between trials when memory cues were presented at different phases of ongoing TWs. Our main hypothesis was that the subject’s reaction time during retrieval would vary according to the instantaneous phase of a cluster’s TW when the memory cue was presented.

In each trial of this task, subjects viewed a list of to-be-remembered items, followed by a retrieval cue. They responded by pressing a button to indicate whether the cue was present in the just-presented list (Fig. 5A, see *Methods*). We measured the instantaneous phase of TWs while they were responding to each cue and tested for a correlation between the phase at each moment and the subject’s subsequent reaction time.

**Figure 5:**
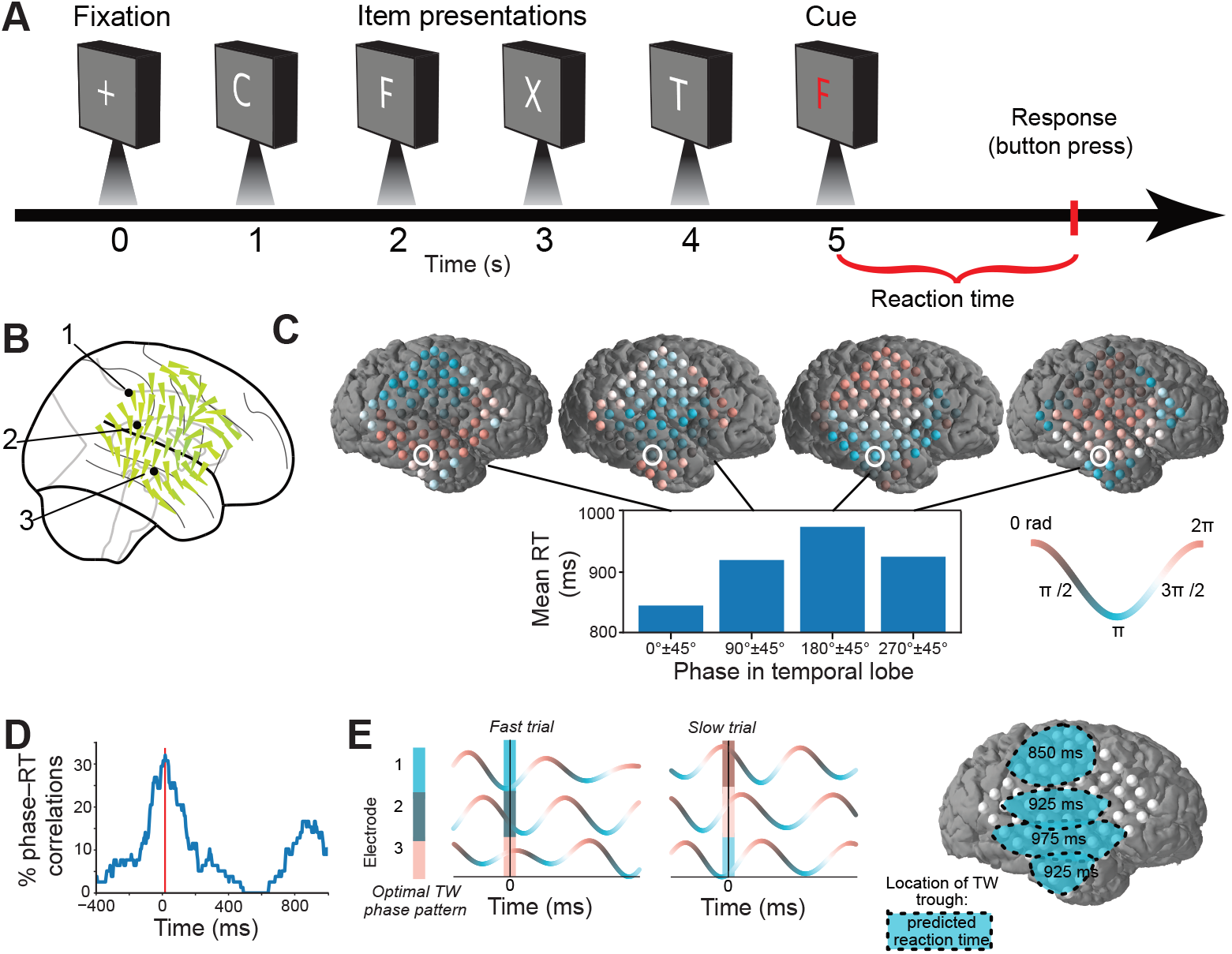
Relation between traveling wave (TW) phase and reaction time in one oscillation cluster. **(A)** Timeline of one trial of the working-memory task. **(B)** Brain map of a 12.5-Hz TW in subject 72, with arrows indicating mean propagation direction across trials. **(C)** Illustration of the link between TW phase and reaction time. Bar chart indicates the mean reaction time computed as a function of the instantaneous TW phase at the moment of cue presentation, as measured on electrode #3 (white circle). Brain plots indicate the topography each of four ±45° phase patterns where electrode color indicates the mean phase. **(D)** The percentage of electrodes in this cluster showing a significant correlation between TW phase and reaction time for each timepoint relative to cue presentation (t = 0). Red line indicates timepoint of peak correlation. **(E)** Schematics illustrating how fast and slow reactions correlated with different TW phase patterns at the moment of stimulus presentation. The phase patterns for fast and slow reactions are shown across three example electrodes in the left panel. Analogously, in the right panel each ellipse indicates how the subject’s reaction time shifts with the position of the TW trough at cue presentation.

The TWs on many oscillation clusters showed a significant correlation between the phase when an item was presented and the subject’s reaction time. An example of this pattern is shown in Figure 5, which shows data from an oscillation cluster where TWs often propagated in a dorsal-to-ventral direction at 12.5 Hz across the right hemisphere (Fig. 5B). Figure 5C plots the mean reaction time of this subject as a function of the instantaneous phase of this TW in the temporal lobe when the item was presented (see also Supp. Video S3. The subject responded fastest (∼840 ms) on trials when the memory cue was presented while the peak of the TW was positioned in the temporal lobe (and, accordingly, when the TW trough was in the parietal lobe). Inversely, the subject responded more slowly (∼975 ms) when the cue was presented while the TW had the opposite phase (i.e., when the TW’s peak and trough phases were located in the parietal and temporal lobes, respectively). This link between the TW phase and efficiency of memory retrieval was robust, as ∼30% electrodes in this cluster showed a significant correlation between the TW phase at cue presentation and the subsequent reaction time (*p*′*s <* 0.05, circular–linear correlation; Fig. 5D). Thus, during memory retrieval this subject’s reaction time varied with the instantaneous phase of ongoing TWs in the right temporo-parietal cortex, with fastest responses occurring if the cue was presented when the peak phase of the TW was in the temporal lobe (Fig. 5E,F).

Other subjects also showed similar patterns, by exhibiting significant links between reaction time in memory retrieval and the phases of ongoing TWs in the alpha band (Figs. 6). Across our dataset, we found a significant link between TW phase and memory retrieval in 23% of all clusters with TWs, including oscillations in both the theta (26%) and alpha (20%) bands (*p*′*s <* 0.05, binomial tests). Together, these results suggest a role for the phase of cortical TWs in coordinating the timing of memory retrieval.

**Figure 6:**
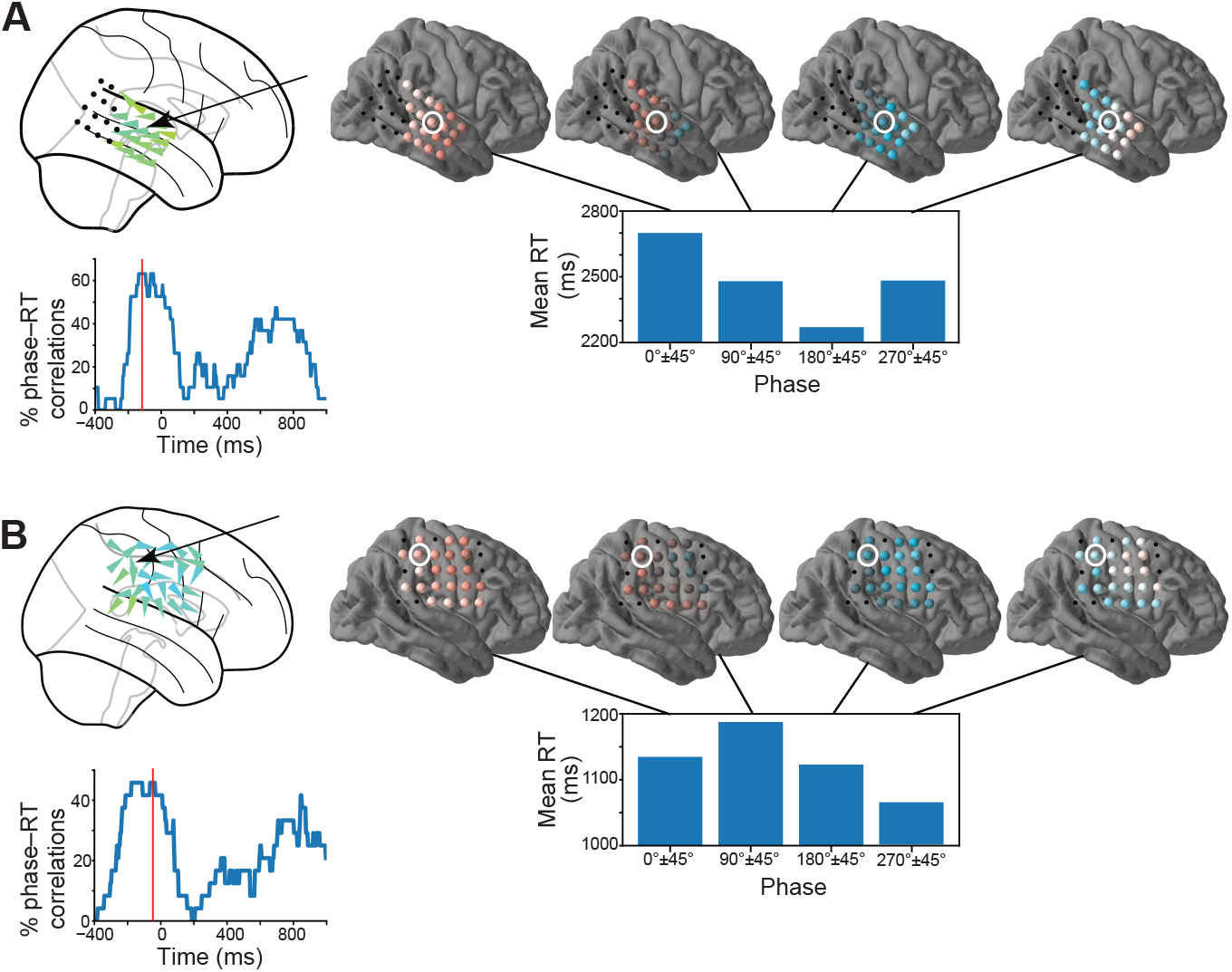
Oscillation clusters showing links between TW phase and reaction time. **(A)** Oscillation cluster showing a 10-Hz TW in subject 73. Red line indicates the timepoint of the peak correlation between TW phase and reaction time. Brain maps following format of Fig. 5. **(B)** Oscillation cluster in subject 46 with a 9.5-Hz TW that exhibited a significant correlation between phase and reaction time that peaked ∼80 ms before cue presentation.

### Timecourse of the link between TW phase and memory retrieval

In the examples shown above, for alpha-band oscillations the correlation between TW phase and reaction time was strongest at the moment when the memory cue was presented. This suggests that the functional role of TW phase in memory retrieval relates to early-stage processes. We next measured the timing of the link between TW phase and reaction time across the dataset. For each electrode in a cluster showing reliable TWs with unidirectional propagation during working memory retrieval, we measured the correlation between reaction time and TW phase throughout cue presentation. We then measured the timepoint of the peak correlation for each cluster. Consistent with the examples shown above, alpha-band TWs generally showed the strongest correlations between phase and reaction time prior to cue presentation (Fig. 7A). This result indicates that the phase of alpha-band TWs correlates with early stages of cue processing.

**Figure 7:**
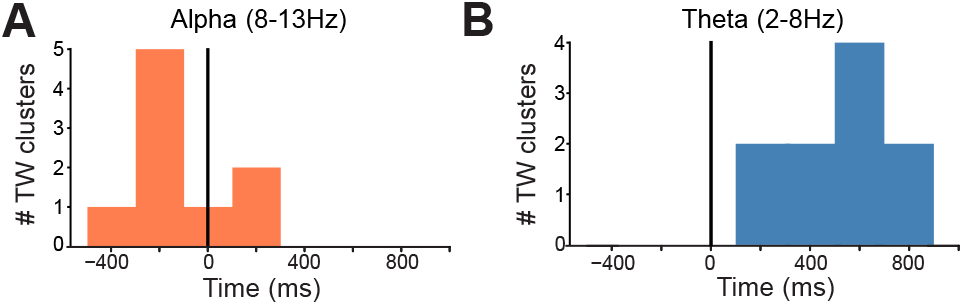
Timecourse of the peak correlation between phase and reaction time for traveling waves at different frequencies. **(A)** For traveling waves in the alpha frequency band (8–13 Hz), histogram shows the timing of the peak link between TW phase and reaction time, time measured relative to cue presentation at 0 ms. **(B)** Same as A for TWs at theta frequencies (2–8 Hz).

In addition to alpha-band TWs, we also found that slower, theta-band TWs showed significant correlations between phase and reaction time. However, this effect occurred ∼600 ms after cue presentation (Fig. 7B), which is significantly later than the timing of the effect for alpha-band TWs (*p <* 10^*−*3^, rank-sum test). This timing shift suggests that alpha- and theta-band TWs have different functional roles in memory, supporting early and late-stage processes, respectively.

## Discussion

A persistent question over the past decades has been how widespread areas of the brain organize their interactions to support different behaviors. TWs provide one answer to this question by propagating in particular directions across the brain to coordinate neuronal activity with high temporal precision. Here we found that the TW direction and timing correlate with memory encoding and recall, which suggests that propagating neural oscillations support cognition by organizing the spatiotemporal structure of neural activity.

Prior studies showed that the theta and alpha oscillations that comprise TWs are phase locked to neuronal spiking and high-frequency oscillations via the phenomenon of phase–amplitude coupling^9,40,41^. With our findings, this suggests that the propagation of theta and alpha oscillations across the brain as TWs indicates when and where the brain is exhibiting discrete pulses, or “packets,” of neuronal activity moving across the cortex^42^. Thus, the propagation direction and phase of theta and alpha TWs may reveal the sequence and timing of when neural representations are communicated across brain regions. These findings have fundamental implications for explaining how different brain regions represent information and interact to support behavior^43^. Our findings also have translational and clinical applications because they suggest that measuring TWs could improve our ability to interface with the brain and diagnose neurological disorders.

A key aspect of our results is identifying a link between distinct directions of TW propagation and separate functional processes, in particular memory encoding and recall. In conjunction with earlier research^25,44,45^, this suggests that a fundamental way in which the brain’s functional connectivity transiently reorganizes is by changing the directional interactions between different brain regions. Because posterior-to-anterior TWs were associated with successful memory encoding and anterior-to-posterior TWs were associated with memory recall (Fig. 8A), it suggests that forming new episodic memories involves the flow of neural activity from posterior regions into the frontal lobe^46^, while retrieval involves the flow of neural activity in the opposite direction^47,48^.

**Figure 8:**
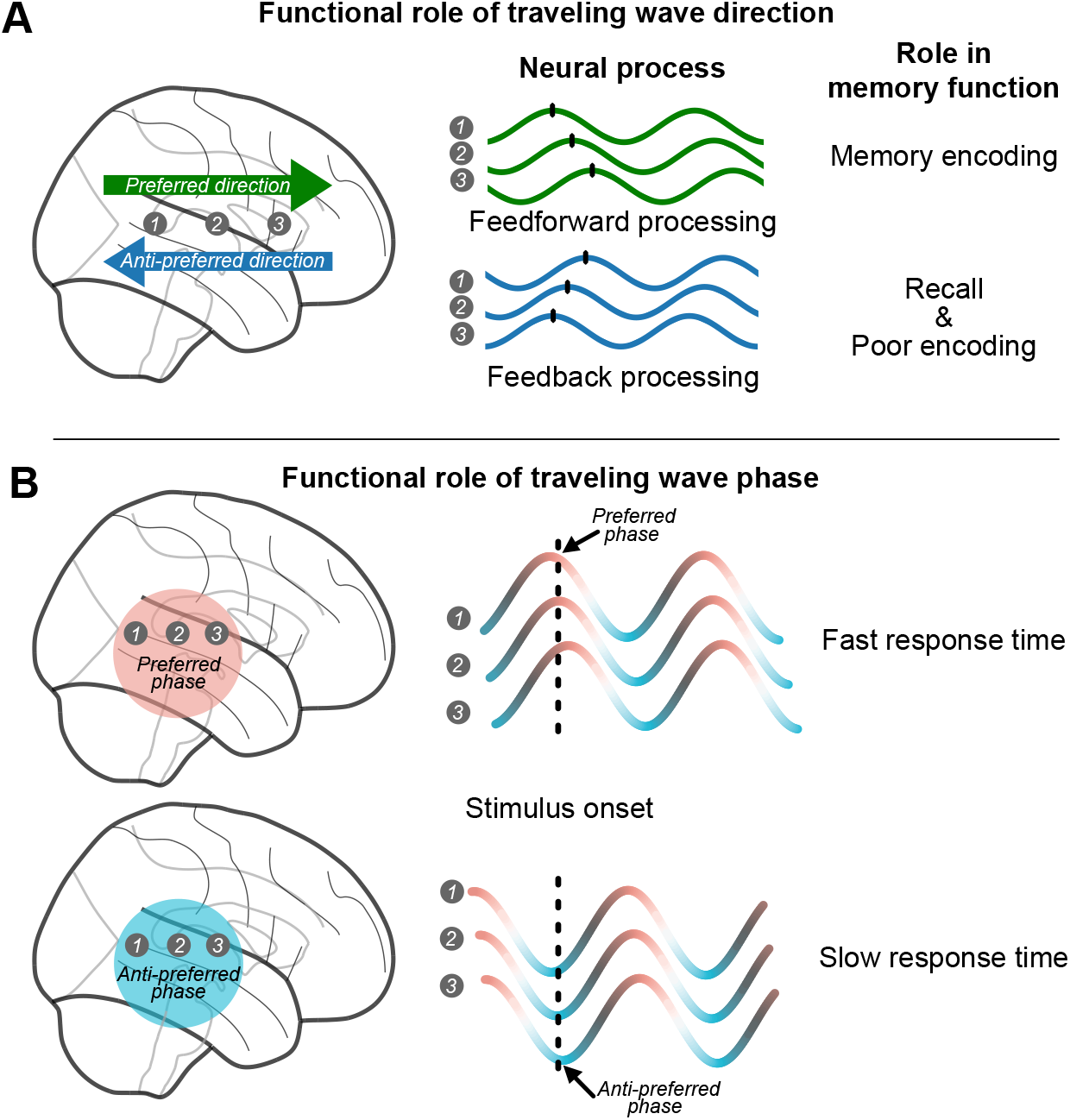
Hypothesized relations between traveling wave (TW) features and memory processes. **(A)** When presented with a list of words during an episodic memory task, successful memory encoding more likely when waves propagated in the preferred direction, as opposed to the anti-preferred direction. We hypothesize that preferred and anti-preferred TW propagation may reflect more general neural processes including feedforward and feedbackward cortical processing, respectively. **(B)** During the retrieval portion of the working-memory task, subjects responded more quickly to memory cues that were presented when the preferred phase of a TW was in a particular region. This suggests that a TW’s phase at each moment indicates whether the brain is primed for memory processing.

One more general possibility is that posterior-to-anterior TWs correspond to feedforward processing while anterior-to-posterior TWs correspond to feedback processing^49–53^. This interpretation is also consistent with earlier work showing that different patterns of neuronal oscillations modulate feedforward networks during visual perception^51,54^ and as well as feedback processing during top-down control and prediction^55^. Consistent with our results, there is also other evidence of neural activity changing direction for specific functional states^25,44,56–60^, thus suggesting that our results are part of a broader phenomenon.

An important question going forward is to understand the mechanisms underlying cortical TWs and, in particular, how TW propagation may shift to support different behaviors. Some work suggests that TWs in the cortex are driven by underlying corticothalamic networks^61,62^ (but see Halgren et al.^63^). Thus, one potential mechanism by which the direction of TW propagation could change is by local increases in excitation at certain thalamic subregions. This excitation could accelerate the frequency of cortical oscillations^20^ and alter TW propagation direction, as predicted by coupled-oscillator models of TWs^1,4,64^. Computational models of TWs could thus be useful for assessing the potential mechanisms underlying memory-related direction shifts.

A TW propagating in a particular direction may indicate that a region is uniquely engaged in a particular functional process. However, a further possibility is that the neural networks in individual regions simultaneously support multiple directionally organized processes, such as concurrent feedback and feedforward processing^50^. Following this view, the propagation direction of TWs at each moment may be informative about the current weighting, or attention, given to each process. Consistent with this idea, prior work demonstrated a link between the amplitude of neuronal oscillations and the attention given to specific neuronal representations^65,66^. In the context of our results, the presence of posterior-to-anterior TWs during successful memory encoding may indicate that the brain is currently attending to feedforward processing to represent the current stimulus and transfer it to memory (Fig. 8A). Inversely, the bidirectional patterns during unsuccessful encoding may indicate that feedforward processes were attended more weakly^67^. Following this logic, the increases in anterior-to-posterior TWs before recall may correlate with top-down processing related to memory search^47,48^.

Our findings suggest that many TWs linked to behavior are endogenous and ongoing in the brain, rather than being evoked by task events. This is most notable for alpha-band TWs, whose direction and phase correlated with performance before stimulus presentation in both encoding and retrieval, indicating that the oscillations were present prior to stimulus onset. This heightened relevance of alpha-band TWs prior to stimulus onset indicates their role in priming relevant brain regions to be in an optimal state for an upcoming visual stimulus^26,27,29,68–70^. In contrast to our alpha-band results at early timepoints, it is notable that we found that theta-band TWs correlated with behavior at later timepoints because it suggests that theta TWs have a fundamentally different functional role^60,71–73^.

In addition to propagation direction, we found that the timing of TW propagation over specific regions was predictive of the speed of memory retrieval. This finding may improve our understanding of earlier studies that showed functional roles for specific phases of ongoing neuFollowing the list, the subjects were presenterdonal oscillations^74,75^, with the propagation of TWs perhaps reflecting moving “packets” of cortical activity to support memory encoding^42,76^. Our results are also consistent with the classic hypothesis that a fundamental function of alpha oscillations is to perform a “scan” for relevant behavioral information^77^. This theory suggested that TWs play a role in the neural basis of attention, with the phase of a TW indicating the specific region of cortex that is preferentially attended at each moment during perception (Fig. 8B). Our findings, with recent work^5^, generally support this TW scanning theory, and our work suggests that this same mechanism may be relevant more broadly to memory^68,78,79^.

It might considered surprising that some of our results were not observed previously, given that human brain oscillations have been measured for decades. It is possible that many previous studies that reported direction- and phase-like patterns in a range of behaviors were actually related to TWs^72,80–82^. Our results relied on new analytical methods, which may have been essential for our findings. In particular, one challenging aspect of measuring TWs in humans is that there is substantial variation in oscillation frequencies and propagation directions across subjects and brain regions. Our analysis framework accommodated this diversity by measuring each subject’s TWs in a customized manner rather than assuming identical propagation and frequencies across all subjects. Given that we observed substantial variability across individuals, it emphasizes the importance of analyzing human brain data in a manner that accounts for intersubject differences in electrophysiology^83–85^. In light of the analytical challenges of measuring TWs in humans and the hints of similar patterns in prior literature, it suggests that TWs may actually have a much broader role in behavior and cognition than previously appreciated.

Traveling waves may be useful for practical purposes, beyond fundamental research. For brain– computer interfacing, TWs might be a useful neural signal for more effectively decoding the brain’s current state. In particular, our direction results indicate that measuring TW propagation can indicate whether the current brain state is well suited for memory encoding. Going forward, it may be possible to use TWs to measure more advanced aspects of cognition, perhaps with the use of improved recording methods, including high-density neural recording arrays^44,86,87^, as well as with noninvasive methods^16,88,89^. Further, TWs could provide biomarkers for identifying neurological disorders related to abnormal neural connectivity such as autism^90^ or epilepsy^91^. Thus, characterizing the directional propagation of TWs holds the potential for new approaches for brain–computer interfacing and disease diagnosis by revealing when the brain’s current communication state is abnormal. TWs may also be useful for guiding the clinical use of brain stimulation, by providing a new target biomarker that reflects neural connectivity.

## Methods

### Participants

The 145 subjects who contributed data to our study were pharmacoresistant epilepsy patients surgically implanted with grids and strips of electrodes on the surface of their cortex for the purpose of identifying epileptogenic regions. The patients’ clinical teams determined electrode place-ment to best monitor each patient’s epilepsy. 68 patients performed an episodic-memory task, and 77 patients performed a working-memory task. Data for the episodic memory task were collected at 8 hospitals: Thomas Jefferson University Hospital (Philadelphia, PA); University of Texas Southwestern Medical Center (Dallas, TX); Emory University Hospital (Atlanta, GA); Dartmouth–Hitchcock Medical Center (Lebanon, NH); Hospital of the University of Pennsylvania (Philadelphia, PA); Mayo Clinic (Rochester, MN); National Institutes of Health (Bethesda, MD); and Columbia University Hospital (New York, NY). Data for the working-memory task were collected at 4 hospitals: Thomas Jefferson University Hospital (Philadelphia, PA); University of Pennsylvania Hospital (Philadelphia, PA); Children’s Hospital of Philadelphia (Philadelpha, PA), and University Hospital Freiburg (Freiburg, Germany). Following approved institutional-review-board protocols at each hospital, all patients provided informed consent.

### Verbal Memory Task

In the episodic memory task, subjects performed a verbal free recall paradigm^31^, in which they were asked to memorize a list of 12 words sequentially presented as text on the computer screen. Figure 2A presents the timeline of an example list. Each word was presented for 1600 ms, followed by a blank screen for 750–1000 ms. Lists consisted of high-frequency nouns (http://memory.psych.upenn.edu/Word_Pools). Following the list, the subjects were presented with a 20-s math distractor task prior to recall. During recall, subjects were given 30 s to verbally recall the words in any order. We recorded the verbal responses on a microphone and then manually scored the recordings after the task.

### Working memory task

Patients performed the Sternberg working memory task^32^. Each trial of the task consisted of two phases. First, during memory encoding, patients were first shown a fixation cross for 1 second with a jitter of ±100 ms followed by a list of 4 items, each presented for 700 ms seconds with a 275–350-ms interstimulus interval. The lists were composed of only consonants to prevent patients using mnemonic strategies such as creating sequences of letters that sound like words. Next, during the memory retrieval portion of each trial, viewed a memory probe item and were instructed to press a key to indicate whether the item was present on the list. The task then indicated whether the patient had responded correctly. Patients performed this task with a mean accuracy of 90% and a median reaction time of 1.16 s.

### Electrocorticographic brain recordings and referencing

During the tasks, data was recorded at 500, 1000, or 1600 Hz using a clinical intracranial electroencephalographic recording system at each hospital (Nihon Kohden EEG-1200, Natus XLTek EMU 128, Natus Quantum EEG, or Grass Aura-LTM64 systems). Subdural grid and strip electrodes had a distance of 10 mm between contacts. Each electrode’s signal was initially referenced to a common contact placed intracranially, on the scalp, or on the mastoid process. We filtered electrical line noise using a 4^th^-order Butterworth notch filter at 58–62 Hz (USA hospitals) or 48–52 Hz (Freiburg). We identified the location of each electrode by co-registering a structural magnetic resonance image (MRI) taken prior to surgery with a computed tomography (CT) image after electrodes were surgically implanted in order to compute electrode locations in standardized Talairach coordinates^92^.

### Identifying traveling waves

We defined a traveling wave (TW) as a single oscillation at one frequency that appears across a region of cortex with a progressive phase shift. To identify TWs in our data, first we used an algorithm to identify spatially clustered groups of electrodes, “oscillation clusters,” that showed oscillations at approximately the same frequency. We then measured whether the phase across these clusters showed the progressive phase shift that characterizes traveling waves^4^. To find these oscillation clusters, we first identified the groups of at least 5 neighboring surface electrodes that showed narrowband oscillations within a 2-Hz window, while being within 25 mm of at least one other electrode with a similar frequency peak. We found the frequency of these narrowband oscillations on each electrode individually by identifying peaks in the power spectrum, which we measured at 200 frequencies logarithmically spaced from 2 to 40 Hz using Morlet wavelets.

Next, building upon methods from Zhang et al.^4^, we identified traveling waves by identifying local plane waves across the electrodes in each oscillation cluster using a circular–linear regression model^34^. To measure the instantaneous phase at each electrode, we first applied a Butterworth filter to each electrode’s signal on each trial, with a filter bandwidth that extended ±15% around the electrode’s mean narrowband frequency. We then measured the instantaneous phase of each electrode’s filtered signal using the Hilbert transform. At each timepoint, we converted the phase at each electrode to a relative phase shift by subtracting, at each timepoint, the mean phase of the oscillations measured across all electrodes in the oscillation cluster. We used circular statistics to manipulate all phase values with the PyCircStat toolbox^93^.

### Measuring local propagation direction

Having computed the relative phase shift on each electrode at each timepoint, we next tested for spatial propagation of the oscillation across the cluster. Whereas our earlier work performed this task by fitting one propagation direction for the entire cluster^4^, instead here we separately fit the direction for each electrode individually. By allowing each electrode to have its own propagation direction, this method had improved sensitivity to TWs with curved propagation patterns, as well as to TWs that were present at only a subset of the electrodes in the cluster.

We fit the circular–linear model for each electrode individually, based on the phase gradient measured on the nearby electrodes (within 25 mm) in the cluster. We only fit this model for electrodes with at least 3 nearby contacts. This procedure measured the features of the TW propagation around each electrode, by quantifying the propagation direction (an angle between *α* ∈ [0^*°*^, 360^*°*^]) and the spatial frequency (*ξ* ∈ [0^*°*^*/mm*, 18^*°*^*/mm*]). To compute these parameters that describe the local TW at each electrode *i*, and timepoint, we fit the equation

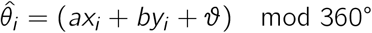

where *a* = *ξ* cos(*α*), *b* = *ξ* sin(*α*), and *x* and *y* are the electrode’s spatial coordinates. Following earlier work^4,34^, we used a grid search to optimize the values for *a* and *b*. This grid search identified the propagation direction and spatial frequency for each TW by minimizing the difference between the predicted phase 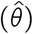 and actual (*θ*) phase values across the nearby electrodes. We measured the statistical reliability of each model fit by computing the circular correlation coefficient between the predicted and actual phases and then adjusting for the number of fitted parameters and datapoints (*ρ*_*adj*_)^4,94^.

Based on applying this model to each electrode individually, we then used two criteria to label an electrode cluster as exhibiting a significant TW on a given trial. First, we required that each cluster have a reliable phase gradient at the group level, as determined by averaging the adjusted correlation coefficient from all the electrodes in the cluster and ensuring it was above 0.2 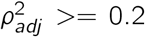 Second, we ensured that the mean power spectrum across all electrodes exhibited a robust narrowband peak. See Fig. S9 for examples of trials without significant TWs. Based on these criteria, we included in our analyses oscillation clusters that had reliable TWs on at least 30 encoding trials.

### Categorization of cluster directionality

Across oscillation clusters, we found TWs that exhibited wide-ranging propagation patterns, including unimodal, bimodal, and multimodal distributions of directions. To characterize these diverse patterns, we designed a method to quantify multimodal directional distributions, rather than only unimodal direction distributions.

To characterize these varying types of propagation patterns, we fit a mixture of von Mises distributions^34^ (the circular analogue to Gaussian distributions) to the distribution of propagation directions from all encoding trials (Supp. Fig. S3). We fit this pattern using a nonparametric model-fitting procedure for circular data, which modeled the overall direction distribution as a mixture of multiple von Mises distributions, each with a different angle and magnitude. In this model, each individual fitted von Mises distribution reflects one particular direction in which the TWs on the cluster frequently propagate. Distributions fitted with more than one von Mises distribution thus showed multiple distinct propagation directions. We used an iterative method to determine the best fitting mixture of von Mises curves, as the sum of the minimum number of von Mises curves (each centered at a different direction) that would fit 99% of the variance in the original distribution of propagation directions^57,95^. We then labeled each cluster as showing unidirectional or bidirectional propagation based on the directions and magnitudes of the mixture of individual fitted von Mises curves. If at least 80% of a cluster’s propagation directions were fit by a single von Mises curve, then we labeled it as showing unidirectional propagation. Likewise, we labeled a cluster as bidirectional if two von Mises distributions (each representing 20–80% of TW directions) were required to capture its propagation distribution. We labeled a cluster as showing “nondirectional” TW propagation if it exhibited no consistent direction over trials (Rayleigh test, *p >* 0.05) or if its propagation patterns could only be accurately fit by a mixture of 3 or more von Mises distributions (this was required in 6% of all clusters).

### Determining a cluster’s preferred propagation direction

Next, for the clusters with bidirectional TW propagation, we tested whether one of the two predominant directions was preferred for memory encoding. To do this, we followed the earlier fitting approach, but applied it just to the trials where memory encoding was successful. We labeled the cluster’s preferred direction as the angle of the von Mises distribution from the overall model fit that was closest to the most prominent propagation direction fit to the successful encoding trials. We determined the preferred angle from the model fit to all trials because this larger dataset provided more precision in categorizing propagation directions as either preferred or anti-preferred. Based on these calculations, we then used the fitted angles to label whether a TW on each individual trial propagated in a direction closer to the cluster’s preferred or anti-preferred direction (Fig. 3C).

### Calculating the relation between TW direction and memory

To measure the timing of the link between a cluster’s propagation direction and memory encoding, we measured the prevalence of TWs moving in the “preferred” versus “anti-preferred” directions at different time offsets relative to stimulus presentation. We performed this calculation separately for trials where the word was successfully encoded as well as for trials where it was unsuccessfully encoded. We determined the cluster’s preferred propagation direction based on the timepoint with the strongest difference in propagation direction between successful and unsuccessful memory encoding, and then recalculated the entire timecourse (2.6 s starting and ending 0.5 s before and after word presentation) of difference scores for each cluster based on that identified preferred direction. We used permutation tests to determine the statistical significance of the relation between TW propagation and memory encoding (see below).

For memory recall, we used a related method to identify the behavioral role of TW direction. At each timepoint relative to word recall, we calculated the percentage of trials with TWs propagating in the cluster’s preferred direction, as determined during encoding. We calculated this for the 3 s prior to word recall or from time of previously spoken word if within 3 s of each other. Because we wanted to measure task-related changes, and individual clusters showed variability in their overall level of TW propagation, we performed a baseline normalization for each cluster. For each cluster, we normalized the observed percent of TWs propagating in the preferred direction relative to the cluster’s non-memory baseline. This baseline included task periods with no stimuli on screen including intertrial intervals.

To examine whether TWs moved in specific anatomical directions for particular memory processes (Fig. 4), across all clusters we computed a weighted distribution of the anatomical directions of TW propagation for each memory process. The weighting for each cluster’s preferred or anti-preferred direction was determined from the percent of individual trials that was captured by that direction’s underlying von Mises curve.

### Measuring the relationship between TW phase and reaction time

To determine the relation between the phase of TWs and reaction time during the retrieval portion of our working memory task, we first calculated the instantaneous phase for each timepoint, on the electrodes in clusters with unidirectional TWs. We focused this analysis on unidirectional TWs to distinguish any observed effects from those related to directional shifts. Then, we used a circular–linear regression to test for a relation between the instantaneous phase at each timepoint relative to cue presentation and the subject’s subsequent (log-transformed) reaction time. Prior to performing this analysis, we excluded trials where a subject showed poor performance. We identified these trials by normalizing each log reaction time to a *z* score relative to the distribution of reaction times in each session and excluding trials with *z* scores above 2. We labeled a cluster as showing a significant correlation between phase and reaction time if it contained at least 10 electrodes in the cluster and at least 25% of those electrodes showed a significant correlation between phase and RT at any timepoint.

### Statistical procedures

We used a permutation procedure to assess whether the directional patterns that distinguished successful versus unsuccessful memory encoding were statistically reliable. We generated 100 random surrogate datasets by shuffling the labels that indicated whether each item presentation was successfully remembered or forgotten. Then, for each random surrogate datasets, we recomputed the entire statistical procedure. This provided a distribution of difference scores that indicated the magnitude of the shift between preferred and anti-preferred propagation directions for successful encoding that would be expected by chance. We tested the significance of the original directional difference scores by comparing its values with the distribution of difference scores from the surrogate data. A difference score was labeled significant if it exceeded the 95^th^ percentile of values from the surrogate distribution (i.e., *p <* 0.05).

For recall, we tested the statistical significance of pre-retrieval direction shifts using two-sided binomial tests. The tests compared the prevalence of preferred and anti-preferred propagation at each timepoint before recall relative to the level in the baseline period for that cluster, correcting for multiple comparisons with the false-discovery-rate procedure^96^.

To test the reliability of memory-related direction changes across all subjects, we used a non-parametric permutation test^97^. This method identified contiguous time periods where TWs showed reliable increases or decreases in preferred or anti-preferred propagation, relative to the timing of particular behavioral events. This procedure assessed significance at the group level for consecutive temporal intervals by comparing the results with those found from applying the same procedure to 1000 surrogate values from random shuffling, with correction for multiple comparisons.

## Acknowledgements

We thank Anup Das, Tom Donoghue, Bard Ermentrout, Molly Hermiller, Lukas Kunz, Serra Favila, Jacqueline Gottlieb, Bradley Lega, Salman Qasim, & Erfan Zabeh for providing helpful critical feedback on the manuscript. We thank Michael Kahana, Paul Wanda, & Joseph Rudoler for providing data and technical support.

## Funding

This work was supported by the DARPA Restoring Active Memory (RAM) program (Cooperative Agreement N66001-14-2-4032) and National Institutes of Health Grants R01-MH104606, U01-NS113198, and RF1-MH114276. The views, opinions and/or findings expressed are those of the author and should not be interpreted as representing the official views or policies of the Department of Defense or the U.S. Government.

## Author contributions

U.M., H.Z., and J.J. designed and implemented the data analyses, and U.M. and J.J. wrote the manuscript.

## Supplemental Information

**Video S1: Example traveling wave (TW) on a trial where memory encoding was successful**. *Animation of a TW in patient 34 during successful encoding (Related to Figure 2). Animation includes filtered signals during trial, local directions indicated on brain map, single arrow of mean direction across electrodes, and topography of TWs phase over time. Single arrow indicating mean direction across electrodes is visible when wave is reliable*.

**Video S2: Example traveling wave (TW) on a trial where memory encoding was unsuccessful**. *Data from patient 34. Same format as Supp. Video S1*.

**Video S3: Animation of a traveling wave that showed a correlation between phase and reaction time during memory retrieval**.

**Table S1:** Prevalence of traveling waves by brain region. The numbers in the left column of this table indicate the total number of subjects with surface grid and strip electrodes in each region. For each combination of region and frequency range, we measured the number of clusters with TWs and then categorized them by their directional patterns. Rows denote combinations of brain region and oscillatory frequency range, separately during memory encoding and recall. Columns denote the category of each TW clusters.

| Region | Patients | Surface Electrodes | Electrodes in oscillation clusters | Oscillatory Range | # clusters | # clusters with TWs | # nondirectional clusters | # unidirectional clusters | # bidirectional clusters | # memory related bidirectional clusters |
| --- | --- | --- | --- | --- | --- | --- | --- | --- | --- | --- |
| **ENCODING** |  |  |  |  |  |  |  |  |  |  |
| **Frontal** | 57 | 1658 | 1391 |  |  |  |  |  |  |  |
|  |  |  |  | **Theta** | 41 | 32 | 3 | 10 | 19 | 14 |
|  |  |  |  | **Alpha** | 12 | 11 | 0 | 4 | 7 | 6 |
|  |  |  |  | **Beta** | 43 | 30 | 0 | 13 | 17 | 10 |
| **Temporal** | 55 | 1351 | 1080 |  |  |  |  |  |  |  |
|  |  |  |  | **Theta** | 22 | 19 | 1 | 10 | 8 | 8 |
|  |  |  |  | **Alpha** | 14 | 13 | 1 | 9 | 3 | 2 |
|  |  |  |  | **Beta** | 28 | 21 | 3 | 12 | 6 | 3 |
| **Parietal/ Occipital** | 57 | 1009 | 935 |  |  |  |  |  |  |  |
|  |  |  |  | **Theta** | 25 | 21 | 1 | 7 | 13 | 9 |
|  |  |  |  | **Alpha** | 10 | 6 | 1 | 3 | 2 | 1 |
|  |  |  |  | **Beta** | 26 | 19 | 2 | 8 | 9 | 5 |
| **RECALL** |  |  |  |  |  |  |  |  |  |  |
| **Frontal** | 58 | 1884 | 1699 |  |  |  |  |  |  |  |
|  |  |  |  | **Theta** | 53 | 44 | 7 | 12 | 25 | 15 |
|  |  |  |  | **Alpha** | 17 | 15 | 1 | 2 | 10 | 5 |
|  |  |  |  | **Beta** | 53 | 48 | 7 | 18 | 24 | 9 |
| **Temporal** | 55 | 1277 | 965 |  |  |  |  |  |  |  |
|  |  |  |  | **Theta** | 19 | 19 | 1 | 9 | 9 | 5 |
|  |  |  |  | **Alpha** | 14 | 13 | 0 | 7 | 6 | 3 |
|  |  |  |  | **Beta** | 23 | 22 | 4 | 8 | 10 | 5 |
| **Parietal/ Occipital** | 59 | 1059 | 992 |  |  |  |  |  |  |  |
|  |  |  |  | **Theta** | 22 | 21 | 2 | 6 | 13 | 7 |
|  |  |  |  | **Alpha** | 17 | 15 | 3 | 4 | 8 | 3 |
|  |  |  |  | **Beta** | 25 | 20 | 5 | 5 | 10 | 4 |

**Table S2:** Subject Table: Verbal Memory Paradigm. Each row summarizes an individual patient’s sex, age, handedness, clinical electrode coverage of surface grid and strip electrodes, frequency and regional properties of clusters with TWs, and memory performance. The “Subject electrode coverage” column denotes the hemisphere, region, and number of surface electrodes in the region. The “Traveling waves” column denotes the hemisphere and region where the majority of electrodes in the cluster were located. The mean frequency of oscillations for the cluster is in parentheses. Region Key: F, Frontal; T, temporal; O, occipital; P, parietal.

| Subject ID | Sex | Age | Handedness | Surface electrode coverage | Traveling waves | Mean recall rate |
| --- | --- | --- | --- | --- | --- | --- |
| 1 | F | 48 | Left | LF (10), LO (5), LP (8), LT (9), RF (9), RO (2), RP (12), RT (11) | RP (7.5), LP (8.0), LP (13.8) | 19.2 |
| 2 | F | 49 | Left | LF (18), LT (3), LT (9), RF (16), RT (12) | LF (19.5), LF (21.3), LF (8.0), RF (19.6) | 39 |
| 3 | F | 39 | Left | LF (50), LO (1), LP (14), LT (27) | LF (18.7), LT (7.8), LF (5.1), LT (12.3), LF (5.7), LF (4.0) | 33.2 |
| 20 | F | 49 | Left | RF (32), RT (2), RO (12), RP (8), RT (29) | RT (14.7), RO (13.6), RT (9.1) | 38 |
| 26 | F | 25 | Unknown | LO (18), LP (29), LT (4) | LP (5.4), LO (18.8), LP (10.6) | 22.7 |
| 32 | F | 20 | Left | LF (6), LP (2), LT (8), RF (4), RP (6), RT (6) | LT (8.1), RP (17.2), RF (7.7), LF (19.2) | 31.7 |
| 33 | F | 32 | Left | LF (6), LP (3), LT (7), RF (8), RO (2), RP (2), RT (12) | RF (22.5), LF (23.0), RT (20.1) | 21.3 |
| 34 | F | 29 | Left | LF (61), LP (4), LT (10), RF (17), RP (9) | LT (8.9), LP (9.3), LF (5.6), LF (17.0), RF (10.1), LF (6.2), LF (12.3), LP (24.5) | 9.1 |
| 36 | M | 49 | Right | LF (20), LT (2), LO (1), LP (22), LT (35) | LT (6.0) | 16.3 |
| 39 | F | 28 | Unknown | RF (63), RP (26), RT (11) | RF (4.7), RF (28.8), RP (7.3) | 22.2 |
| 42 | F | 28 | Unknown | RF (18), RP (32), RT (22) | RP (8.6), RP (18.5), RP (5.2), RP (3.8) | 63 |
| 45 | M | 51 | Left | LF (21), LP (3), LT (11), RF (5), RP (1), RT (6) | LF (7.8), LF (23.1), LF (7.7), LT (17.8), RF (7.7) | 32.7 |
| 50 | M | 20 | Unknown | LF (3), LO (3), LP (35), LT (23) | LP (7.0), LP (16.2) | 42 |
| 51 | F | 25 | Left | LF (15), LP (9), RF (8), RP (14), ROther(2) | LF (19.3), RP (14.2), LF (5.4), RP (3.4) | 39.6 |
| 53 | F | 39 | Unknown | RF (27), RT (1), RP (9), RT (27) | RF (15.5), RF (6.4) | 17.8 |
| 60 | F | 37 | Unknown | RF (30), RP (6), RT (28) | RT (8.2) | 30.4 |
| 66 | M | 39 | Left | LT (2), LO (1), LP (8), LT (11), RF (5), RT (12), RO (1) | RF (7.9) | 33.2 |
| 68 | F | 39 | Left | RT (3), RO (11), RP (17), RT (31) | RT (17.5), RO (15.0), RP (10.5) | 52 |
| 69 | M | 27 | Left | LF (43), LP (37), LT (8) | RF (15.8), RF (4.6), RP (7.4), RF (22.0), RP (5.7) | 28.7 |
| 76 | M | 30 | Unknown | LF (32), RF (2), RP (9) | LF (5.8), LF (19.4), RP (5.7), LF (11.5), RP (16.3) | 25.7 |
| 84 | M | 25 | Left | LF (38), LP (41), LT (4), RF (3) | LF (5.4), LF (21.0), LF (8.5), LP (7.2) | 17.7 |
| 86 | M | 21 | Left | LF (29), LT (2), LO (1), LP (2), LT (39), RF (3) | LF (5.9), LF (20.1) | 30 |
| 89 | M | 36 | Left | LF (2), LT (2), LT (6), RF (26), RT (3), RP (26), RT (37) | RT (7.7), RT (18.9) | 16 |
| 102 | M | 35 | Left | LF (12), LP (6), LT (12), RF (10), RT (11) | LF (20.1), LF (5.9), RF (19.9), RT (3.5), RF (5.8), LP (7.1) | 25.3 |
| 104 | M | 22 | Left | RF (49), RP (21), RT (22) | RF (6.4), RF (26.5), RP (19.3) | 31.3 |
| 112 | F | 30 | Unknown | LF (3), RF (17), RO (1), RP (1), T (24) | RT (6.9), RT (15.4) | 12.7 |
| 128 | M | 26 | Unknown | RF (10), RP (8), RT (6) | RF (6.7) | 47 |
| 129 | F | 35 | Unknown | LF (28), LOther (10), LP (6), LT (3), RF (27), RP (4), Rother (2) | RF (13.5), RF (4.4), LP (13.1), LF (4.9), LF (14.6), LP (4.8) | 17.5 |
| 130 | M | 57 | Left | LF (40), LP (32), RF (8) | LF (5.5), LF (11.0), LF (19.0), RF (5.2), LP (4.2) | 19.6 |
| 135 | M | 48 | Left | RF (15), RO (5), RP (8), RT (7) | RF (23.3), RF (4.5), RF (12.8) | 8.9 |
| 136 | F | 57 | Unknown | LF (23), LT (4), LO (1), LP (7), LT (34), RF (1), RT (18) | LT (8.3), LT (18.7), RT (7.6), RT (16.2) | 19.8 |
| 142 | F | 44 | Unknown | LF (13), RF (47), RP (4), RT (17) | LF (11.0), RF (5.5), RF (11.4), LF (5.4), RT (14.9), RF (10.3) | 16 |
| 147 | M | 48 | Left | LF (32), LO (6), LP (28), LT (41), RF (1) | LT (20.0), LT (6.3), LT (10.1), LP (13.6) | 18.1 |
| 149 | F | 28 | Left | LF (17), LT (11), LP (5), LT (39) | LT (8.6), LT (19.5), LT (4.5), LF (23.1) | 21.3 |
| 151 | M | 36 | Left | LF (9), LP (4), LT (7) | LP (7.6), LP (18.2) | 27.5 |
| 154 | F | 36 | Left | LF (20), LP (4), LT (40) | LT (11.4), LF (22.9), LT (17.8), LT (4.5), LP (21.8) | 30.1 |
| 155 | M | 37 | Left | LF (43), RF (23), RP (18) | LF (5.0), LF (16.1), RP (4.9), RP (14.2), RF (16.8), LF (5.1), LF (4.7) | 27.5 |
| 159 | F | 43 | Left | LF (38), LT (2), LP (14), LT (34), RF (9), RP (2), RT (10) | LF (20.9), LT (7.9), LP (11.1), LF (6.5), LT (15.3), RF (4.9) | 23.8 |
| 162 | F | 31 | Unknown | RF (11), RT (2), RO (3), RP (6), RT (23) | RT (7.7), RT (16.9), RF (5.2), RF (8.2), RF (19.0), RP (5.1) | 25.7 |
| 167 | M | 33 | Left | LF (20), LO (1), LP (4), LT (42), RF (1) | LT (7.9), LT (21.0), LF (7.9), LF (19.0), LP (18.6) | 44.6 |
| 172 | F | 22 | Left | RF (10), RP (2), RT (12) | RF (22.6) | 19.3 |
| 175 | M | 34 | Unknown | LT (16), RF (29), RO (5), RP (22), RT (18) | RF (6.0), LT (6.2), RP (14.5), RF (18.9), LT (6.1), RT (21.6), LT (14.1) | 22.7 |
| 184 | M | 42 | Unknown | RF (3), RO (2), RP (27), RT (16) | RP (5.8), RP (14.8), RF (19.5) | 25.5 |
| 186 | M | 28 | Unknown | LP (3), LT (8), RF (32), RP (29), RT (31) | RF (6.2), RF (20.0), RP (14.1), RT (9.3), RF (5.4) | 8.3 |
| 187 | F | 52 | Unknown | LF (49), LT (2), LO (3), LP (30), LT (19), RF (1) | LF (16.1), LP (7.8) | 13 |
| 195 | M | 44 | Left | LF (11), LP (8), LT (7), RF (13), RP (5) | RF (24.3), RF (10.4), LP (22.5), LT (7.8), LF (14.7) | 30.4 |
| 196 | M | 19 | Unknown | LF (30), LO (1), LP (28), LT (11) | LP (7.2), LP (18.9), LT (21.6), LT (7.0) | 37.3 |
| 201 | M | 37 | Left | RF (16), RT (3), RO (5), RP (28), RT (33) | RP (10.7), RP (16.4) | 23 |
| 202 | F | 30 | Unknown | LF (4), RF (14), RO (14), RP (5), RT (26) | RT (10.4), RT (20.3), RF (5.2), RO (15.0) | 18.8 |
| 215 | F | 51 | Left | LF (16), LT (32) | LT (8.0), LT (16.9) | 24.3 |
| 221 | M | 57 | Left | LF (41), LT (39), LP (18), RF (1), RP (1) | LT (14.5), LT (8.9), LF (21.8), LF (16.6) | 30 |
| 222 | F | 21 | Left | LF (50), LP (10), LT (21), RF (11) | LF (19.9), LF (5.6), LF (8.7) | 27.2 |
| 226 | F | 42 | Unknown | LF (31), LP (8), LT (35) | LT (22.5) | 25.6 |
| 232 | M | 28 | Left | LF (13), LP (11), LT (18), RF (10), RP (14), RT (14) | RP (6.0) | 47 |
| 234 | M | 25 | Unknown | LF (2), LP (5), LT (49), RT (24) | LT (13.7), LT (3.9), RT (13.9), LT (22.4), LT (6.2), RT (3.7) | 25.5 |
| 250 | M | 30 | Left | LF (1), LO (4), LP (10), RF (8), RO (23), RP (55), RT (20) | RP (20.2), RP (6.7), RP (12.3), LP (20.5), LP (12.0), LP (6.9), RP (12.8) | 45.7 |
| 260 | F | 57 | Unknown | LF (28), LT (46), LO (4), LP (18) | LT (20.3) | 25.4 |
| 291 | M | 35 | Left | LF (21), LP (23), LT (32) | LT (6.7) | 35.7 |

**Table S3:**
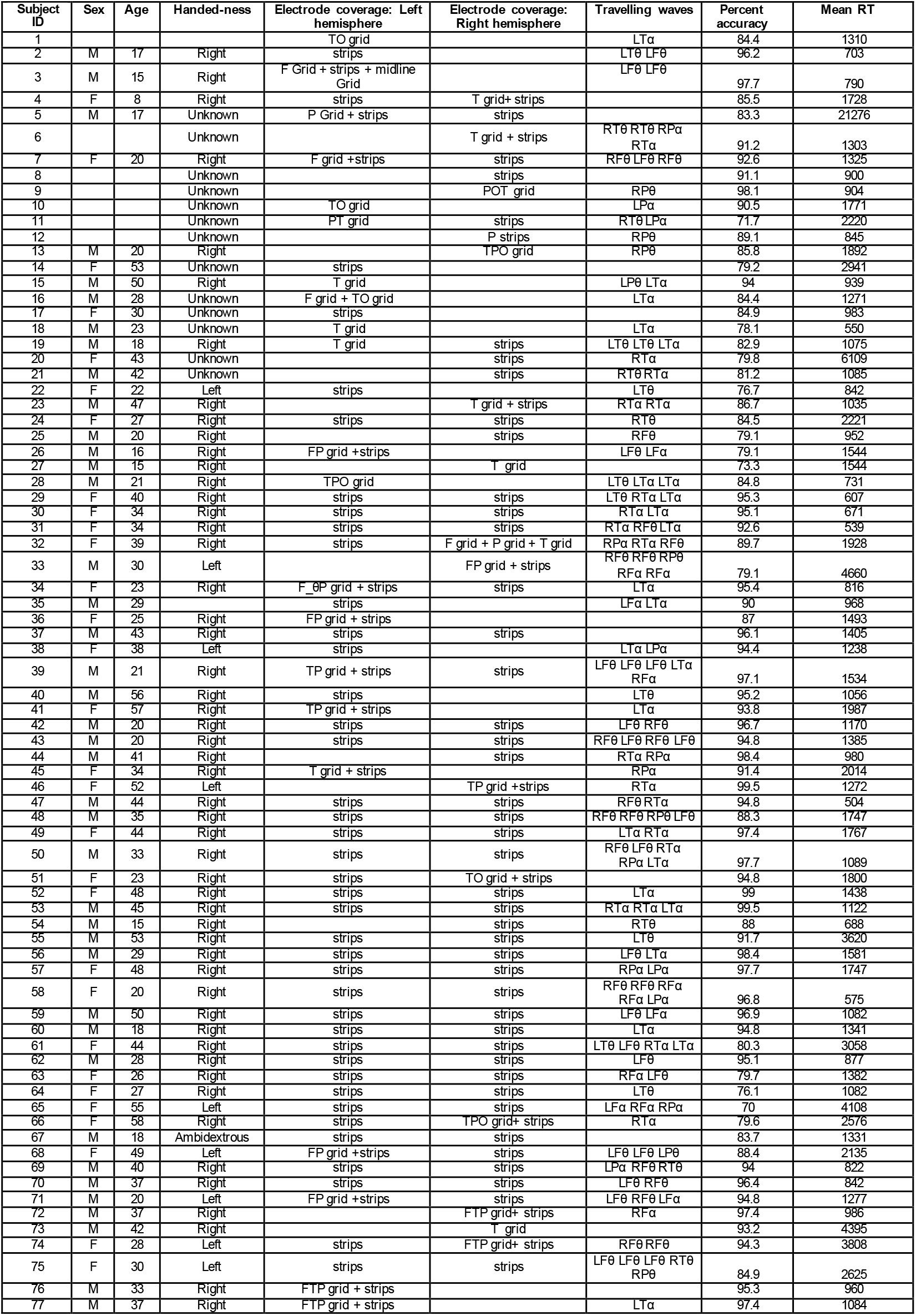
Subject Table: Sternberg Working Memory Task. Each row summarizes an individual patient’s sex, age, handedness, clinical electrode coverage of surface grid and depth electrodes, oscillatory band and regional properties of clusters with TWs, memory performance, and mean response time. The “Electrode coverage” columns denote the regions with suface electrodes and types of electrodes. The “Traveling waves” column denotes the hemisphere, region where the majority of electrodes were located in the cluster, and the band of the peak frequency of the cluster. Region Key: F, Frontal; T, temporal; O, occipital; P, parietal.

**Table S4:**
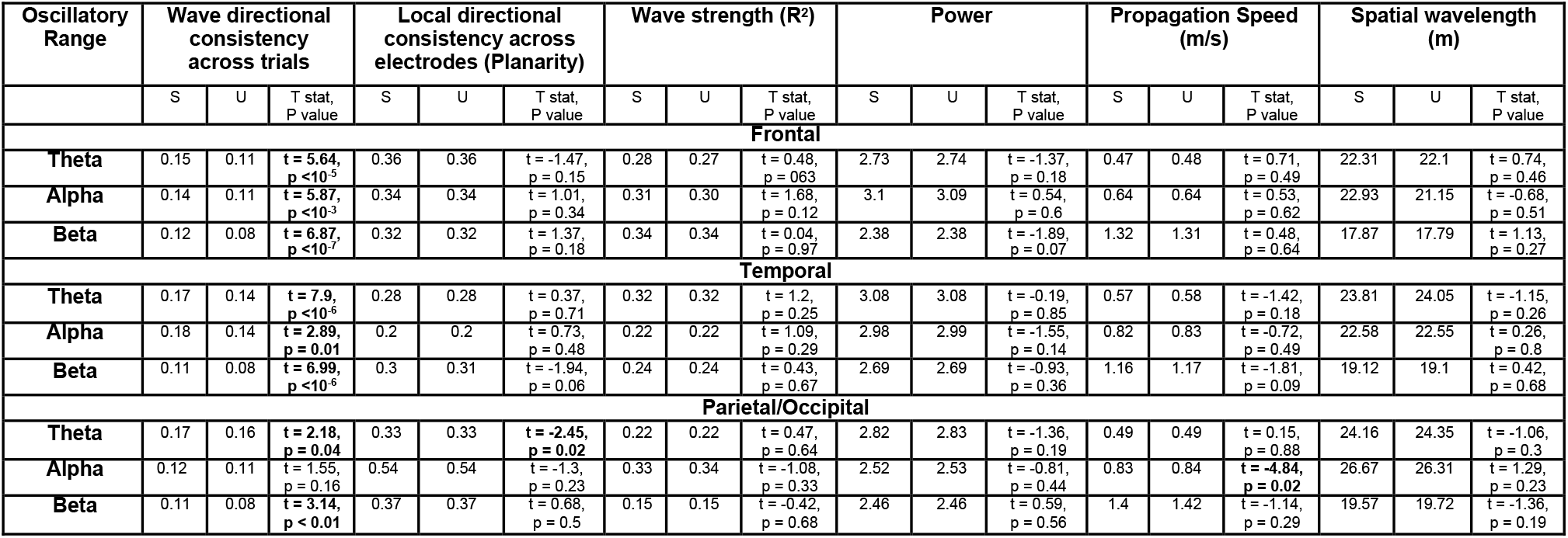
Relation between memory encoding and traveling wave characteristics. Rows denote each combination of region and oscillatory range. Each group of columns shows the relation between memory and the following TW features: wave directional consistency across trials, local directional consistency within a wave (measured across electrodes), wave strength, narrowband power, propagation speed, and spatial wavelength. Within each group, columns “S” and “U” indicate the mean value of each traveling-wave characteristic (see Methods) on trials with successfully and unsuccessfully encoding, respectively. We measured the statistical significance across clusters with TWs with paired t tests.

**Figure S1:**
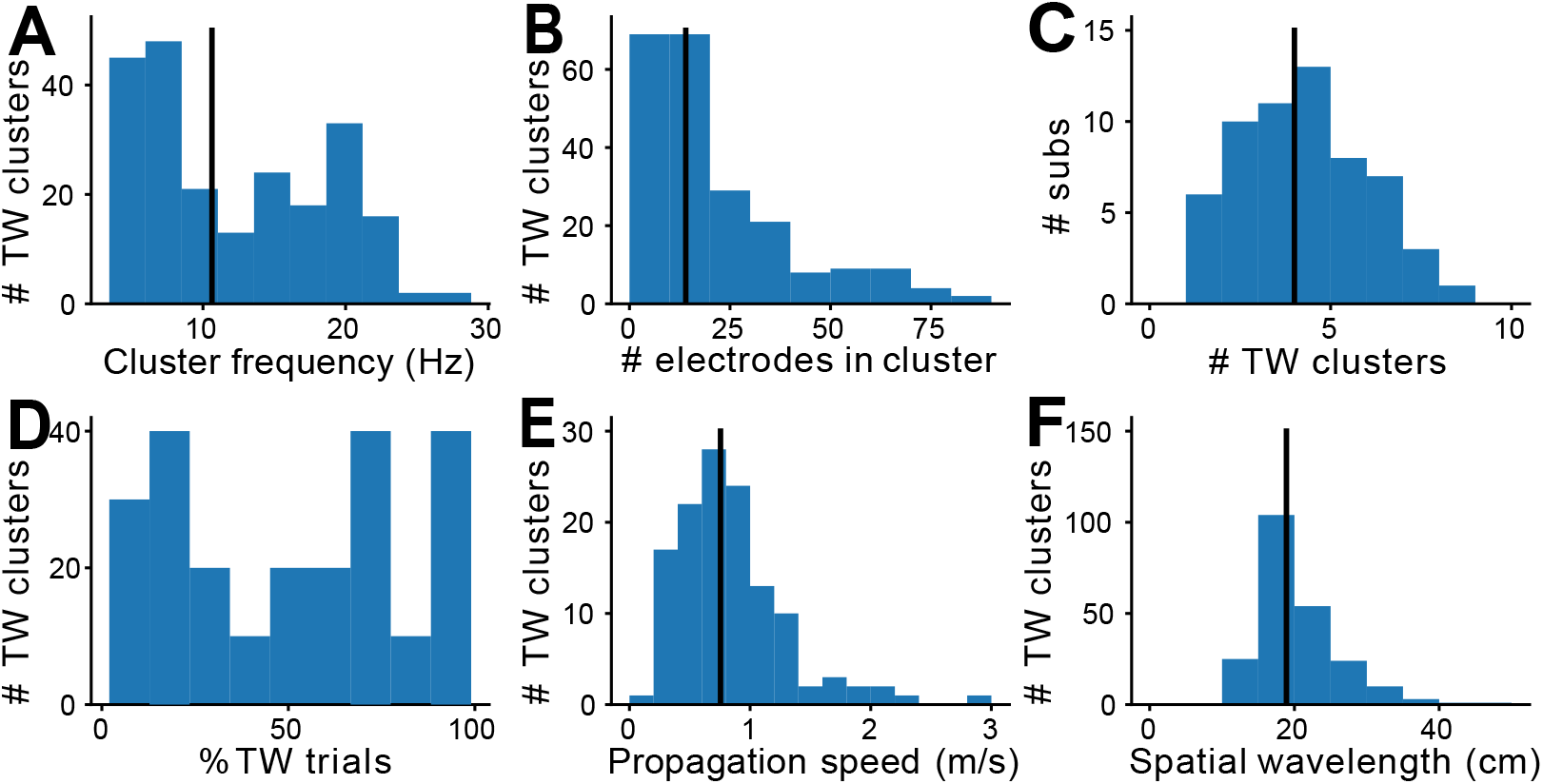
Characteristics of cortical traveling waves in the episodic memory task. **(A)** Histogram of the peak oscillation frequencies for clusters with TWs. **(B)** Histogram of the number of electrodes in each cluster. **(C)** Histogram of the counts of clusters per patient that showed TWs. Most subjects had 2 to 4 clusters across different sets of grid and strip electrodes or groups of electrodes with oscillations at different peak frequencies. A few patients had 5 or more. Patients with many clusters often had multiple smaller clusters of 5-6 electrodes in different regions and hemispheres. **(D)** Distribution of the percentage of single trials that show reliable TWs for individual clusters. **(E)** Histogram of TW propagation speed across clusters. Black line indicates median. **(F)** Histogram of TW spatial wavelength.

**Figure S2:**
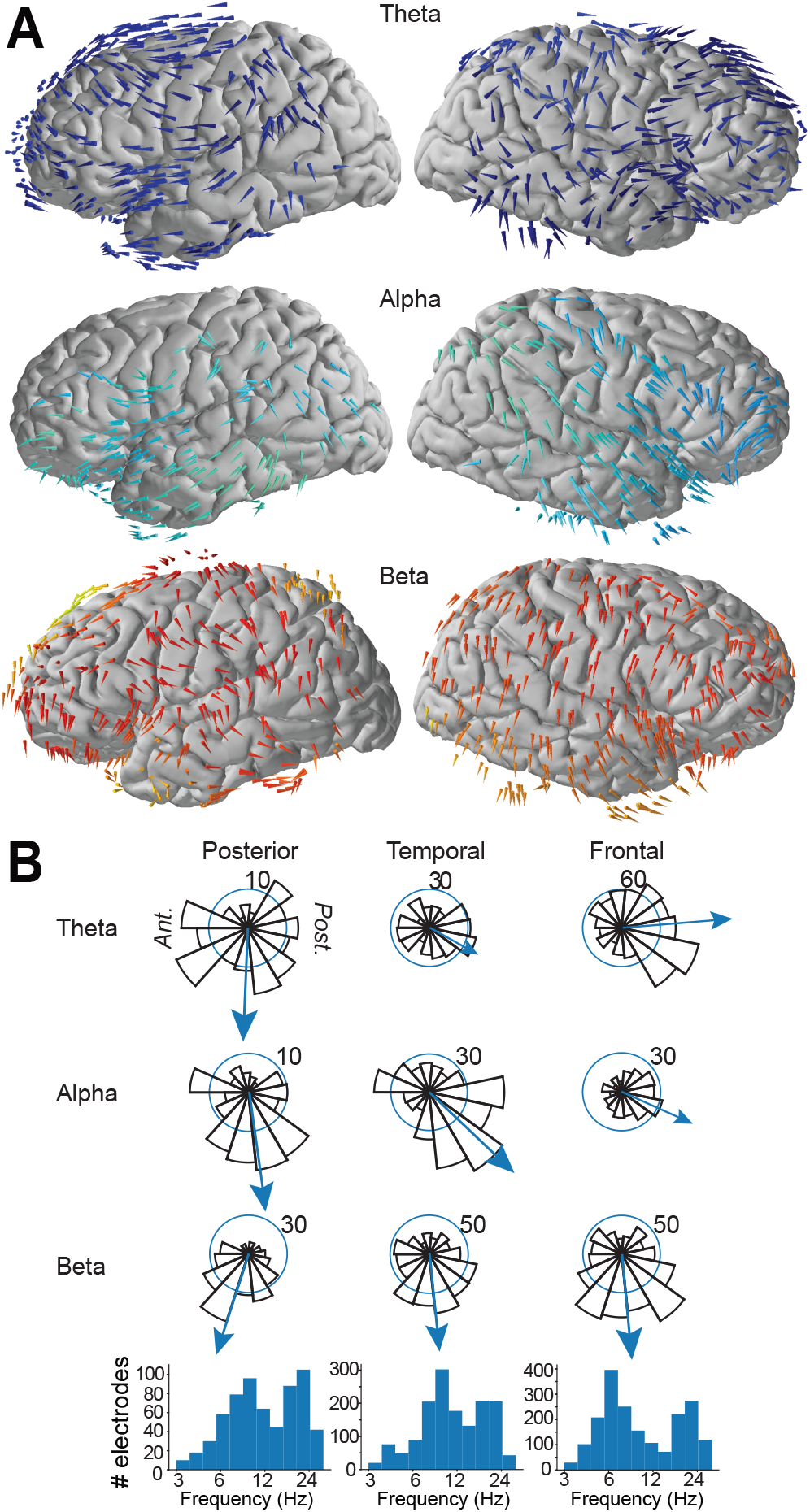
Summary of traveling wave propagation directions in the working memory task. **(A)** Arrows indicate the mean propagation directions of TWs in the theta (blue), alpha (teal), and beta (orange) frequency bands. The density of colored arrows indicates the prevalence of traveling waves at each frequency and the orientation of each arrow indicates the mean propagation direction. **(B)** Distribution of mean propagation directions for oscillation clusters across brain region and oscillatory bands. Bottom: histograms of the peak frequencies of electrodes by cortical region. These plots indicate that Theta-band TWs are common in the frontal and temporal lobes and usually propagate in a posterior-to-anterior direction. Alpha band TWs are present throughout the brain, except for the left frontal lobe. Alpha TWs generally propagate anteriorly, although alpha waves propagated posteriorly in some subjects. This plot shows that traveling waves in the beta band were present throughout the brain; however, they showed substantial variability in propagation directions as compared to the traveling waves seen in the theta and alpha bands.

**Figure S3:**
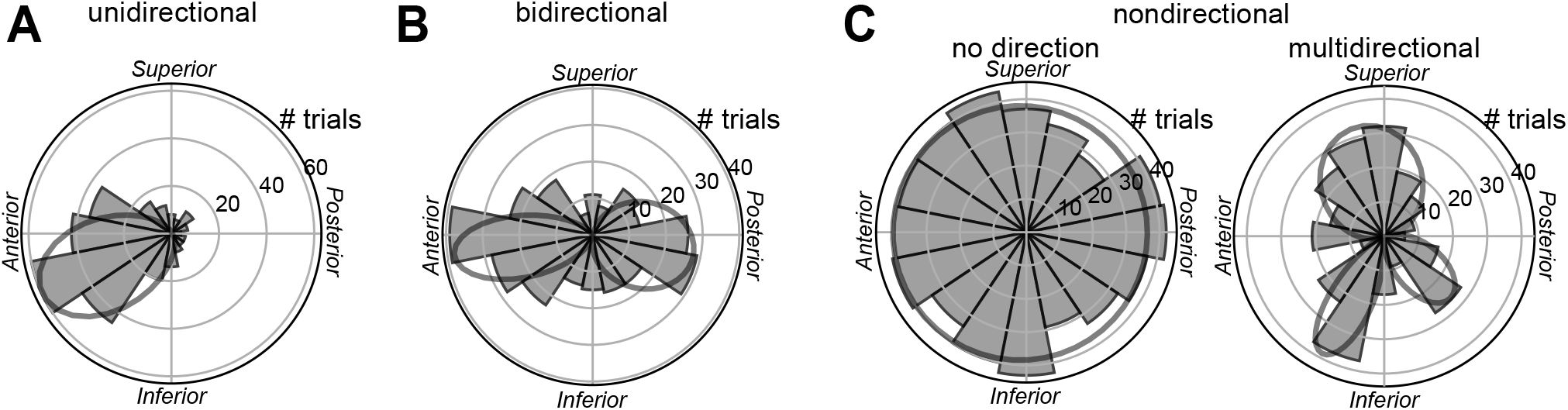
Examples of clusters that showed traveling waves with different types of directional propagation patterns. Plots show example direction distribution for TWs we labeled as propagating in **(A)** unidirectional, **(B)** bidirectional, and **(C)** nondirectional fashions.

**Figure S4:**
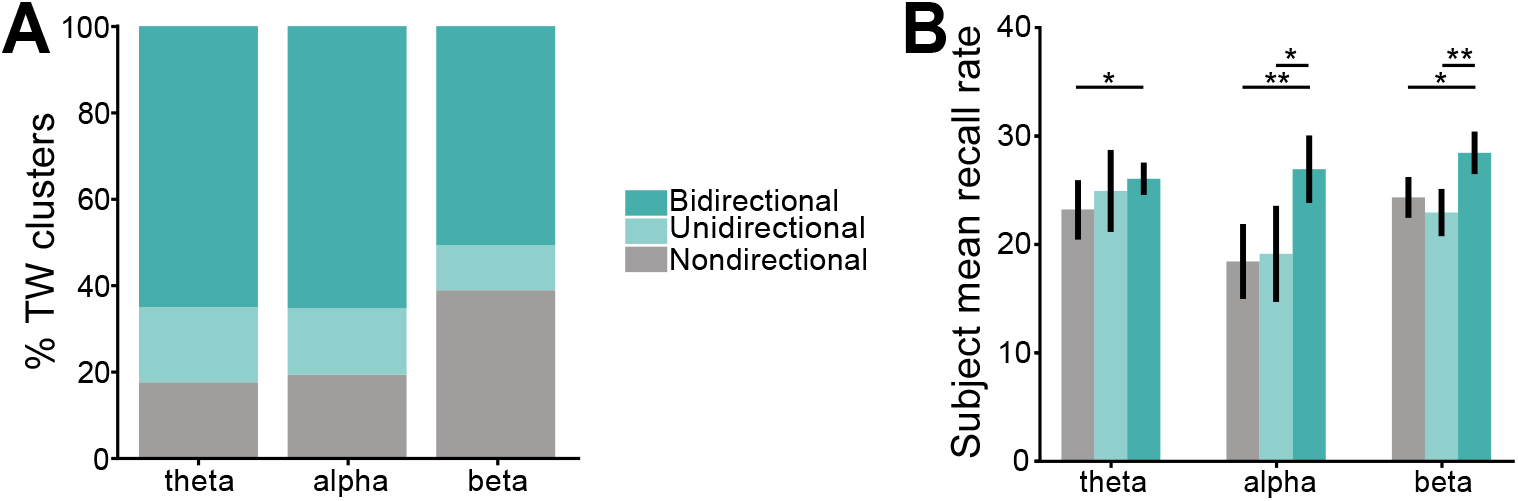
Population categorization of cluster direction patterns in episodic memory task. **(A)** Percent of TW clusters in each oscillatory range identified as bidirectional, unidirectional, and nondirectional. **(B)** Mean percent recall rates for subjects that showed a TW cluster with unidirectional, bidirectional, and nondirectional TW propagation, by frequency band (linear mixed effects model, bidirectional vs. unidirectional clusters: p=0.055; bidirectional vs. nondirectional TW clusters:, p=0.017). Error bars denote ±1 SEM. (∗p < 0.05, ∗ ∗ p < 0.01).

**Figure S5:**
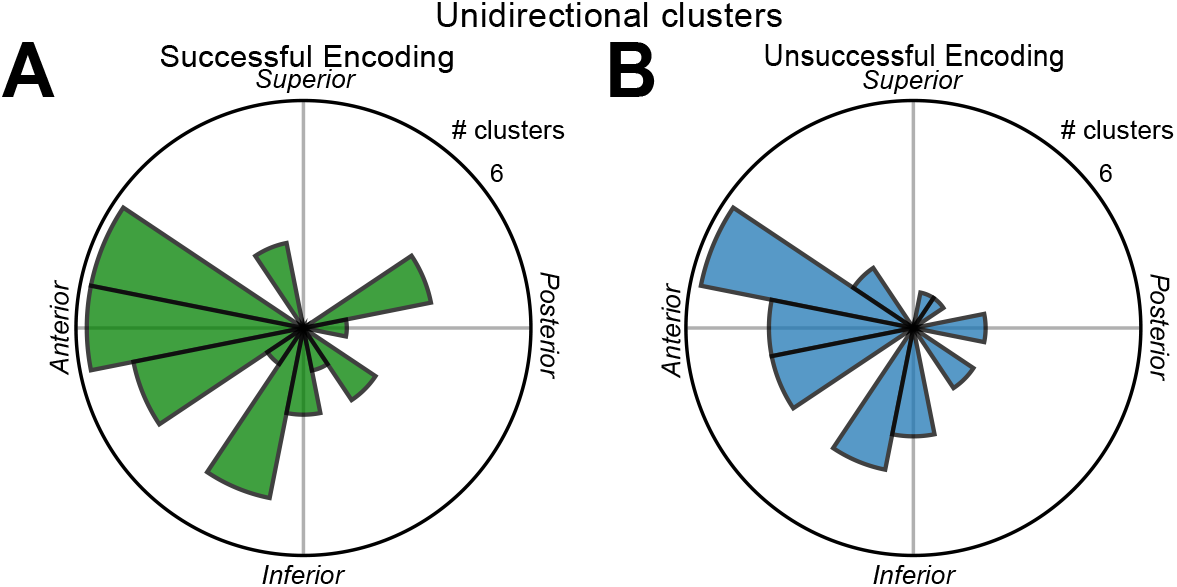
Mean propagation directions during successful (A) and unsuccessful (B) memory encoding for the clusters labeled as unidirectional. A direct comparision showed that the propagation directions of unidirectional clusters did not significantly differ between successful and unsuccessful encoding trials (Non parametric multi-sample test for equal medians, z = 0.91, p = 0.34).

**Figure S6:**
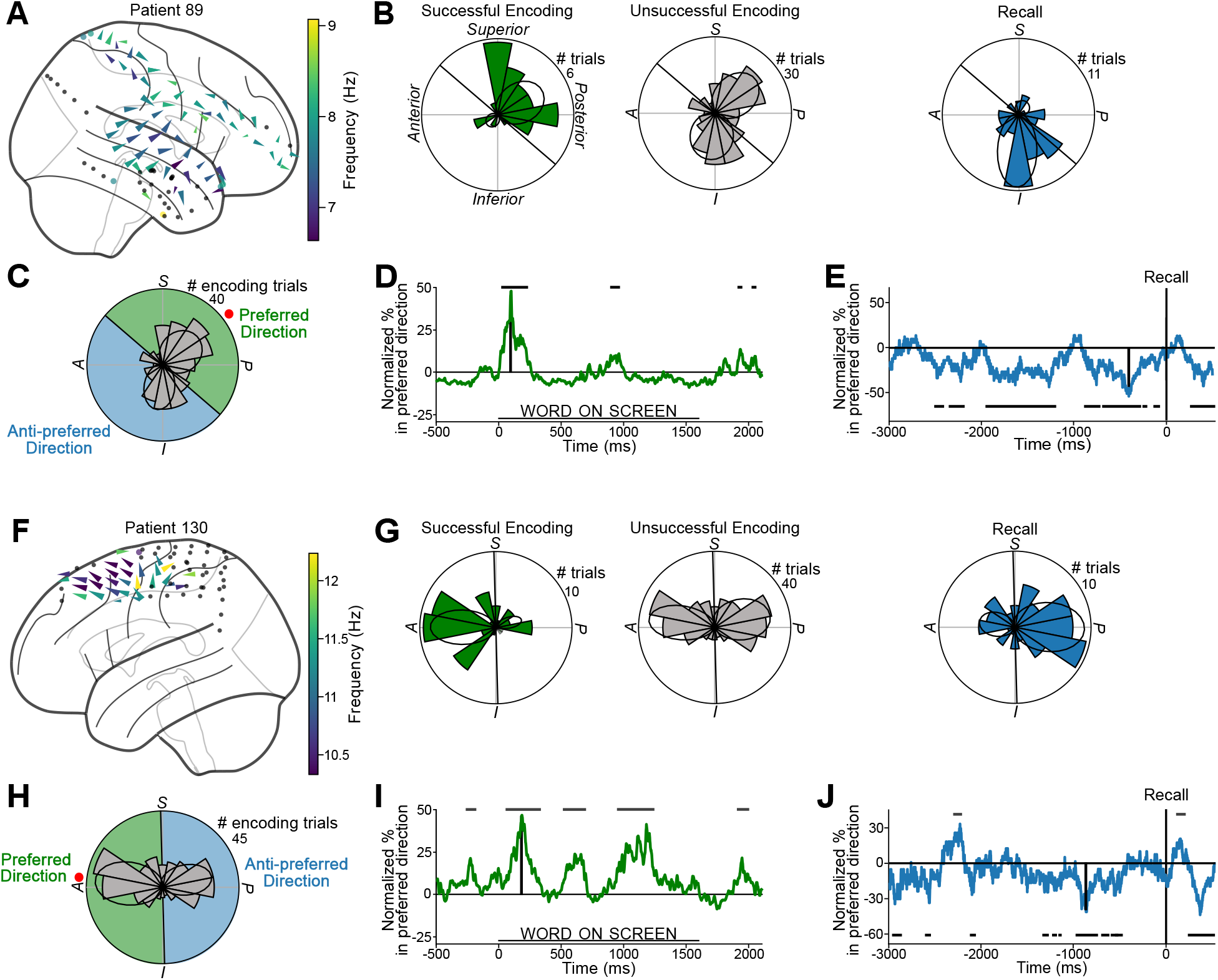
Traveling waves in example subjects who showed a link between TW direction and memory. **(A–E)** Example traveling wave in patient 89 at 7.8 Hz; format of individual plots follows Figure 3. **(F–J)** Example traveling wave frontal cortex of patient 130 at 10.8 Hz.

**Figure S7:**
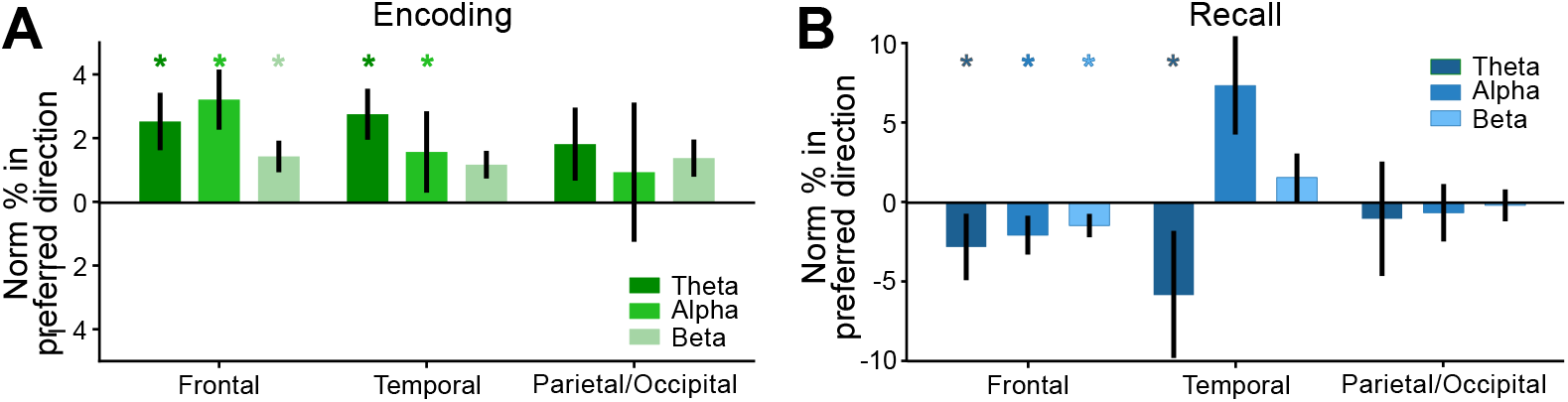
Relation between TW directional shifts and memory processing. **(A)** Normalized difference in the prevalence of TWs propagating in the preferred versus anti-preferred direction for successful relative to unsuccessful memory encoding (averaged across word presentation interval). Asterisks indicate specific regions and oscillatory bands where the normalized percent of TWs traveling in preferred directions across clusters is significantly above or below a distribution of shuffled TW directions (p′s < 0.05, binomial tests). **(B)** Normalized difference of TWs propagating in preferred versus anti-preferred direction averaged during 2 seconds prior to verbal recall. normalized narrowband power

**Figure S8:**
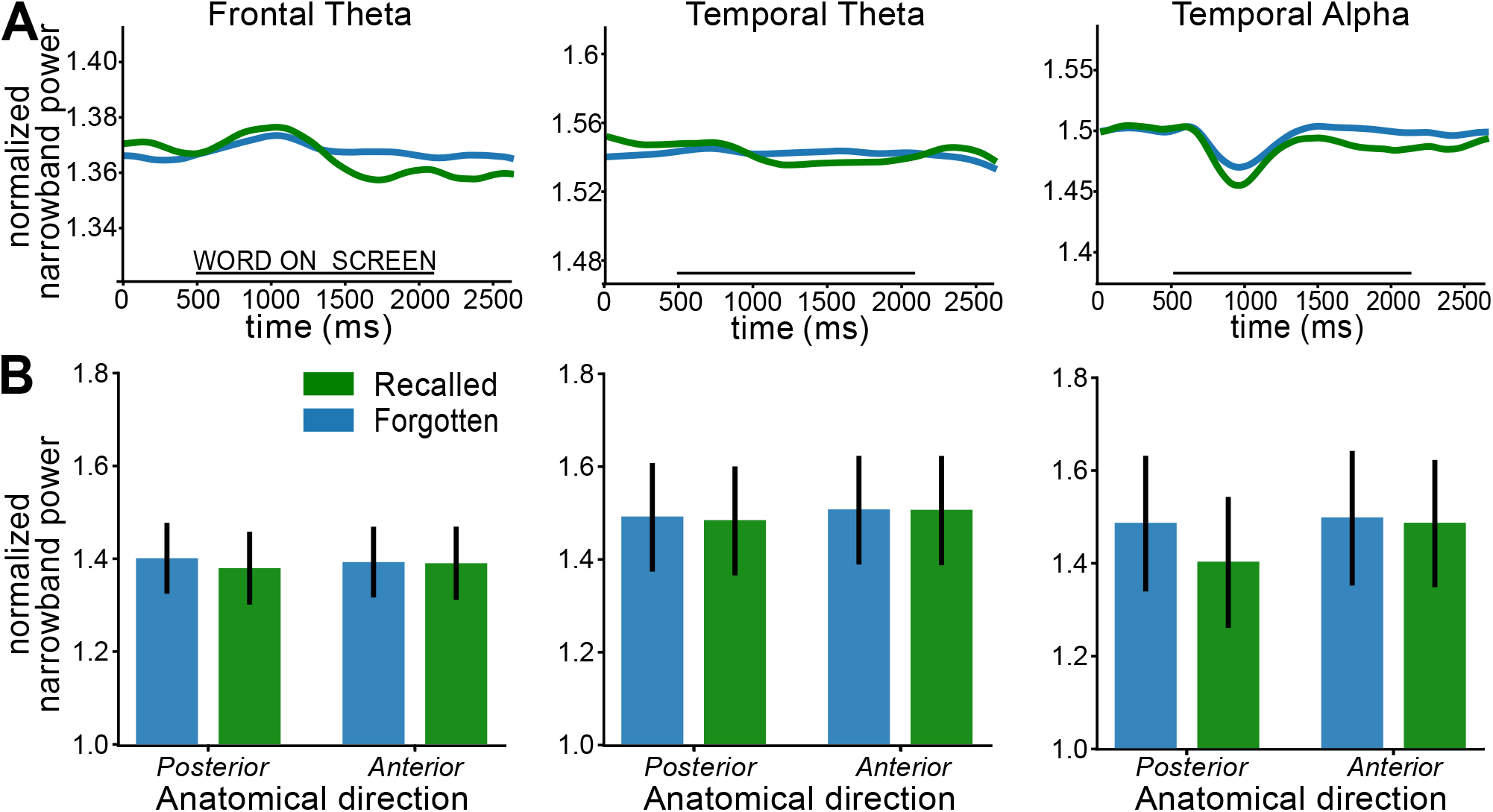
Narrowband power at oscillation clusters that showed traveling waves in the episodic memory task. **(A)** Mean normalized narrowband power centered around each oscillation cluster’s peak frequency, calculated with the log-transformed amplitude of the Hilbert transform. **(B)** Mean normalized narrowband power for oscillation clusters that showed traveling waves averaged over time, separately calculated during time periods when TWs moved posteriorly and anteriorly, during successful and unsuccessful encoding trials. There were no significant differences in mean power across the clusters that showed posterior and anterior propagation (p′s > 0.05).

**Figure S9:**
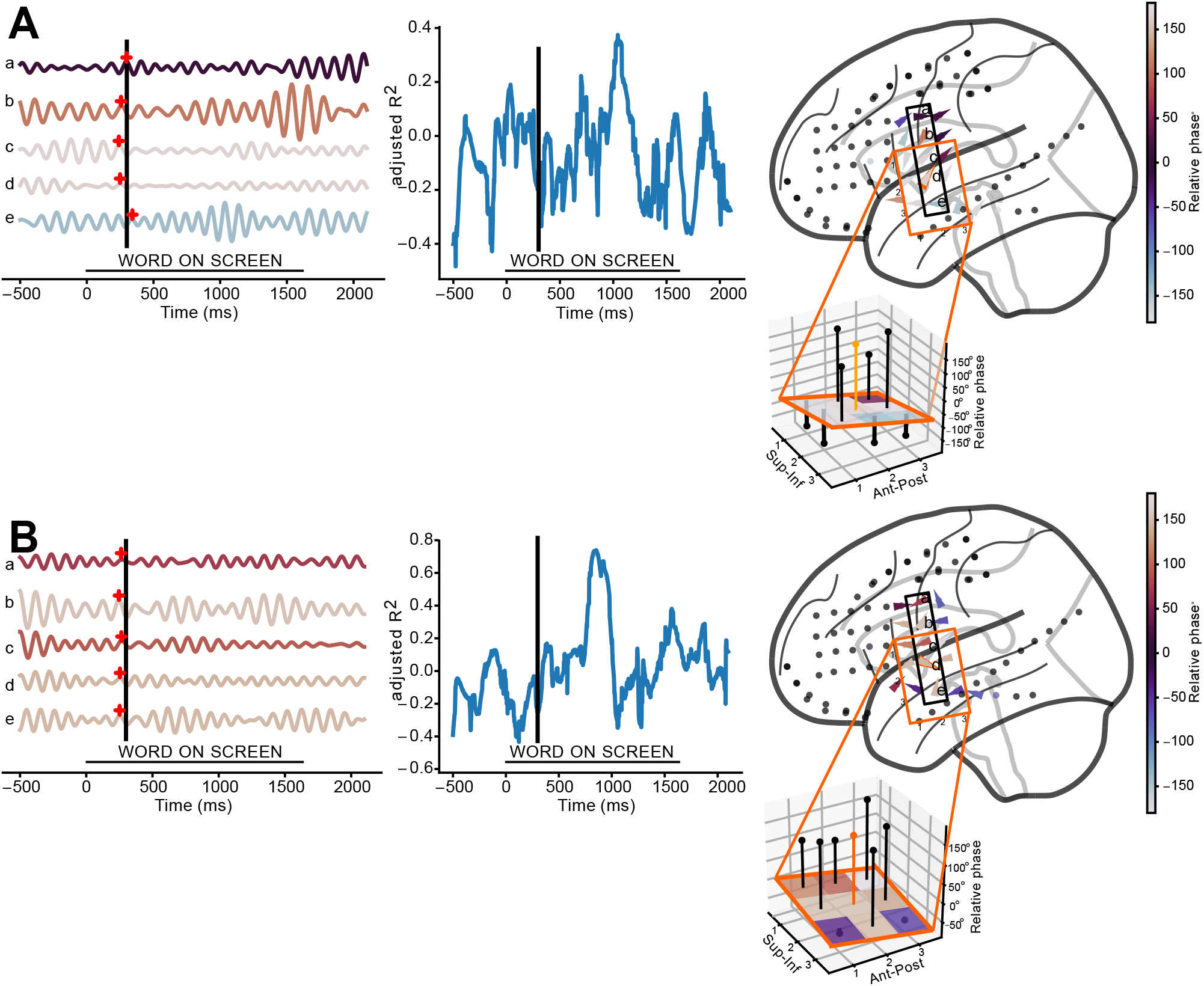
Example data showing the absence of traveling waves. **(A–B)** Example trials where a traveling wave was not present on a cluster that often showed 8.9-Hz oscillations that propagated as TWs on other trials. Data comes from from subject 34 (Fig. 1).

## Notes

### Competing Interest Statement

The authors have declared no competing interest.

